# A climate-driven compartmental model for fungal diseases in fruit orchards: The impacts of climate change on a brown rot-peach system

**DOI:** 10.1101/2022.09.13.507724

**Authors:** Daniele Bevacqua, Chiara Vanalli, Renato Casagrandi, Marino Gatto

## Abstract

As a well-known driving force of diseases in crops, climate change is likely to impact future crop yields. In the present work, we account for climate-related influences on the key parameters of a SIR-type epidemiological model for fungal diseases in stone fruit aimed at simulating different observed epidemic patterns, and, eventually, evaluating the possible impacts of climate change on the disease-induced yield loss. Brown rot disease in peach orchards is used here as a study system. We contrasted simulation results with epidemiological measures collected from an experimental orchard in Avignon (southern France) in two consecutive years, characterized by dissimilar brown rot outbreaks. The capacity of our climate-driven model to adequately reproduce the observed disease patterns suggests temperature and precipitation as key drivers of brown rot epidemics. Overall, the model predicts a potential decrease of brown rot severity under warmer and drier climatic conditions. To comprehensively understand the effect of future climate change on peach yield, alterations of crop phenology must also be accounted for. We thus build a model that considers the synergism between the two factors: vulnerability to the pathogen and varying phenology. Using plausible climate change scenarios, we find that the peach yield in the considered Mediterranean region will be considerably impaired: although brown rot-related yield losses are expected to decrease in warmer and drier climatic conditions, climate change will hinder fruit blooming and consequently the yield because milder winters will impede the achievement of dormancy. By deepening our understanding of climatic impacts on crop fungal infections, the present study may serve as a useful tool to plan and implement suitable adaptation strategies for peach cultivation.

## Introduction

Climate has significant impact on plant diseases, because it potentially modifies host physiology and resistance and it alters pathogen rates of development (Coakley et al., 1999; Juroszek et al., 2020). The variation of climatic variables, due to global warming, has already been proved to have impacted the dynamics of numerous infectious plant diseases and depleted crop production (Bebber, 2015; Romero et al., 2022). At different latitudes, productivity and infection risk are likely to show different responses to climate changes (Chaloner et al., 2021). Abiotic stresses such as heat and drought may not only contribute to increase plants susceptibility to pathogens, but can also alter general defense pathways that disrupt plant resistance (Desaint et al., 2021). Any pathogen can survive only in a given range of temperatures, and experienced temperatures also affect its generation times (Garrett et al., 2006). Similarly, precipitation influences plant disease dynamics by altering both the physiology of plants and the ability of pathogens to survive, disperse and infect hosts (Colhoun, 1973; Milici et al., 2020).

Fungal diseases are responsible for important crop losses worldwide (Fisher et al., 2012) and are a paradigmatic case of climate dependency (Gange et al., 2007; Corredor-Moreno and Saunders, 2020; Romero et al., 2022). For example, spore germination of the rust fungus *Puccinia substriata* has been found to increase with temperature, and the reproduction of root rot pathogens *Monosporascus cannonballus* occurred more quickly at higher temperatures (Garrett et al., 2006). Overall, an increased impact of fungal pathogens due to the lengthening of the growing season by temperature increase has been experimentally proved by Roy et al. (2004) and has been observed in the field (Richerzhagen et al., 2011; Salinari et al., 2006). While lengthening some phases, climate change may also shorten others, like the time duration of susceptible plant phenological stages (Craufurd and Wheeler, 2009). Also, it can disrupt the synchronization between host and pathogens (Marçais and Desprez-Loustau, 2014; Scheffers et al., 2016), so that a potential increase of crop production can in principle be speculated. Water availability also plays a role in infections. Rain induces short-distance spore dispersal and secondary infections through a phenomenon known as “splashing” (Colhoun, 1973; Vidal et al., 2017). Also, infection, corresponding to spore germination and penetration into the host tissue, depends on the surface wetness, which is ultimately affected by precipitation and temperature conditions (Huber and Gillespie, 1992). Although there is empirical evidence provided by lab experiments that fungal diseases require optimal ranges of temperatures and of wetness duration to develop and grow (Phillips, 1982; Biggs et al., 1988; Xu et al., 2001), there is a lack of knowledge of how, and to what extent, many processes simultaneously involved in a fungal epidemic ultimately determine disease dynamics observed in the field (Bebber, 2015). Understanding how climate affects outbreaks of fungal infections is essential to guarantee food security in a context of global change and to plan adaptation actions. These optimized interventions must be aimed to minimize not only the yield losses, due to epidemics, but also the economical and environmental costs related to the disease management (Fisher et al., 2012).

Mathematical models are useful tools to understand the interactions of climate with infectious diseases (Goudriaan and Zadocks, 1995; Savary and Willocquet, 2020; Truscott and Gilligan, 2003). Quantitative modeling approaches are in fact of key importance for assessing the impact of climate change by facilitating the comparison of multiple scenarios (Coakley et al., 1999). This is particularly true for the so-called process-based models, whose parameters have a clear biological meaning and that permit the inclusion of their climatic dependence. These models can then be used to explore crop-disease interactions and to evaluate their possible behaviours in climatic conditions not experienced in the past. The dependence of fungal infections on climatic conditions started being observed around the end of the 20^th^ century, but it is only quite recently that climate-driven epidemiological models have been developed. Magarey et al. (2005) proposed a simple model for predicting periods of infection by fungal foliar pathogens. They modeled the response of the infection rate to temperature and scaled it to the surface wetness duration requirement of the considered pathogen. The model was validated against already published data of several species, including some in the genera *Cercospora, Alternaria, Puccinia*, and then used to evaluate its suitability for risk assessment studies (Bregaglio et al., 2012). Audsley et al. (2005) developed a model estimating green area loss due to winter wheat foliar diseases (leaf blotch, yellow rust, powdery mildew and brown rust), integrating the effect of air temperature, rainfall and relative humidity. In a somehow similar work, Calonnec et al. (2008) proposed a model that considers air temperature, wind speed and direction as climatic inputs affecting parameters of grapevine plant growth and powdery mildew spread (such as the latent period, infection rate, lesion growth, conidial spore production and release). Wang et al. (2021) developed a crop model that (using air temperature, relative humidity, precipitation, wind speed and leaf wetness duration) can assess rice blast impacts. They used it to evaluate consequences of future climate conditions over northern Italy rice production. Even though compartmental SIR-type (Susceptible-Infected-Removed) models have been widely used in plant pathology because of their tractability and extensibility and their plasticity in providing a mechanistic understanding of the epidemic dynamics (Van der Plank, 1963; Cunniffe et al., 2012; Burie et al., 2008), very few studies have specifically investigated how climate conditions can simultaneously affect key epidemiological traits in crop diseases (e.g. pathogen exposure and viability) (Chaloner et al., 2019). Furthermore, including climate forcing in SIR-type models allows the understanding of time-varying pathogen infectivity which, as argued by Cunniffe et al. (2015b), is a current challenge for plant disease epidemiology.

In a recent work, Bevacqua et al. (2018) presented a SIR-type model, more precisely an SEI (Susceptible-Exposed-Infected) model, to describe the temporal spreading of a fungal disease in fruit orchards. The model was parameterized using field data, sampled in the 2014 growing season (*i.e*. from May to July) from a peach (*Prunus persica*) orchard infected by brown rot in Avignon (southern France). Brown rot is caused by *Monilinia* spp. and is one of the most serious diseases of fruit (Villarino et al., 2013; Rungjindamai et al., 2014). The genus *Prunus*, which includes species with important economic value such as plum, peach, apricot, cherry and almond, is severely affected in temperate regions (Lino et al., 2016), with all significant associated economic losses. Brown rot is well known to depend on weather conditions, that influence the fungus life cycle and the overall infection dynamic (Tamm et al., 1995, 1993; Biggs et al., 1988; Phillips, 1982). The parameter values calibrated in the original work presented by Bevacqua et al. (2018) were representative of the average ecological and physiological conditions prevailing in the 2014 growing season. The relevant basic reproduction number evaluated in that study indicated that the environmental conditions met in the field for the 2014 season were extremely favorable to disease development. Although the model closely fit the temporal evolution of the fruit abundance in the different epidemiological compartments in the calibration year, it did not succeed in reproducing the dissimilar epidemic patterns observed in the same orchard for the subsequent growing season (May-July 2015). In other words, using the model with parameter values calibrated on 2014 field data did not succeed in matching the recorded 2015 field data. Since the 2014 and 2015 datasets were collected from the same plants, subjected to the same agricultural practices (*i.e*. irrigation, fertilization, thinning), a candidate cause for such failure was that the model version presented by Bevacqua et al. (2018) did not account explicitly for meteorological conditions, which were indeed quite different in those two years (as it is normal because of inter-annual variability). Taken together, the 2014 and 2015 datasets could however provide an excellent opportunity to evaluate the potential role played by variations of meteorological conditions (therefore climate) in shaping the infection dynamics, opportunity that can be explored by explicitly incorporating their effects on the model parameters.

In this study, we aim at *i*) incorporating the climatic dependence of the epidemiological param-eters in a compartmental model that describes the infection dynamic of brown rot, *ii*) evaluating if this climate-driven model can reproduce the contrasting epidemics observed in two following years *iii*) assessing the climate change impacts on the peach yield losses across the 21^st^ century. First, we built upon the original model proposed by Bevacqua et al. (2018),, testing if the original model was mechanistically unfit to simulate the disease spread under different environmental conditions. Then, because we are considering two different fruit growing seasons in 2014 and 2015, we explicitly added natural fruit abscission to the the original model (Bevacqua et al., 2018), which considered only the 2014 data, thus neglecting possible variation between years..

Next, we accounted for the response of epidemiological model parameters to climatic conditions, contrasting different possible climatic effects and testing different hypotheses to select those that find more support in the field data. We eventually proposed a novel model formulation, explicitly accounting for the effect of climatic data, such as average daily air temperature and precipitation. Those data were selected because those two quantities alone proved already to be able to successfully reproduce very different epidemiological patterns. We then tested the model sensitivity to climatic conditions, altering both daily temperature and precipitation regimes, and we used it to predict the response of a virtual peach-brown rot pathosystem to climate variation observed in the 2010-2019 decade and predicted for the 21^st^ century in Avignon, under different climate change scenarios.

## Materials and methods

### Available data

Data were collected in 2014 and 2015 from an experimental orchard of 43 peach trees (cultivar Magic/GF677) planted in 2006 at the INRAE station of Avignon (southern France, 43°60’ N, 4°49’ E), composed of nine rows of five trees with 4-m spacing between trees (Bevacqua et al., 2018). In the absence of fungicide treatment after bloom, brown rot naturally built up in both years, with higher incidence in 2014. Apart from fungicide treatment, we managed trees according to conventional practices regarding fertilization, irrigation, and pesticide treatments. From the beginning of June until harvest time in mid-July, the abundance of symptomless (see below) and infectious fruits was measured every week of both years on 18 trees after having excluded trees on the border sides and a specimen that displayed very low vigor. Hereinafter, with the term “symptomless”, we refer to fruits that do not present brown rot symptoms following a visual inspection *in situ*.

From the trees on the borders of the experimental orchard, fruit fresh weight was measured from the beginning of May until mid-July in 2014 and 2015. These trees are excluded from the observation of the epidemiological dynamics, because removing fruit could have influenced the spread of the disease by encouraging an unnatural (and additional) dispersal of spores. Over the course of the growing season of 2014 and 2015, 713 and 2,024 total fruits were sampled, respectively (see Figure S1). Blooming time (*t_B_*) and harvest time (*t_H_*) occurred respectively on February 28^th^ and July 15^th^ in 2014 and on March 15^th^ and July 17^th^ in 2015. Fruit abscission is a natural process that varies with fruit load and environmental conditions. The natural fruit abscission rate, *i.e*. the rate at which fruits naturally drop from trees, affects all fruit independent of their epidemiological state. Therefore, we used total fruit abundance (symptomless and infected fruits) to estimate fruit abscission rates (see Table S1 and Figure S2). In addition, we used data published by Xu et al. (2001) to estimate how the mortality rate of the fungal spores depend on temperature conditions. Local data of daily average temperature (°C) and daily precipitation for the period 2010-2019 (see Figure S3) were obtained from the meteorological station of INRAE Saint-Paul (43.92°N; 4.88°E).

We used scenarios (in the form of time series) of future daily average temperatures and precipitation occurrences across the 21st century as they were generated by the project Drias, developed by Meteo-France, with a spatial resolution of 8 × 8 km^2^ (see Figures S6 and S7).

### Model outline

Here, we provided an essential description of the model originally presented and calibrated in Bevacqua et al. (2018). A sketch of it is schematically represented in Figure 1. The model was used to describe the temporal dynamics of the host fruit in the growing season, from the time of pit hardening *t*_0_ to harvest time *t_H_*, in relation to brown rot infection. Fruits can be in one of the following states at time *t*: susceptible (*S*(*t*) or simply *S*), exposed to the pathogen (*E, i. e*. spores are present on the fruit cuticle. This is a necessary, but not sufficient, condition to become infected), or infected and infectious (*I*). The probability that a susceptible fruit is exposed to the pathogen in a unit of time is assumed to be proportional to the density of infectious fruits *I*, via a time-invariant coefficient of proportionality. An exposed fruit can either return to the susceptible state at a time-invariant rate due to spore mortality or it can progress to the infectious state at a rate that is proportional to the fruit fresh weight which, provided that the fruit has attained a critical size (Gibert et al., 2007), is strictly related to the cuticle crack surface area. Fruit fresh weight was assumed to vary in time according to a logistic equation. Infectious fruits are removed from the host population at a constant rate since they dry out and stop being sources of new inoculum. For a more extensive explanation of the rationale justifying the above mentioned assumptions, as well as for the references supporting it, the reader can refer to Bevacqua et al. (2018).

**Figure 1:**
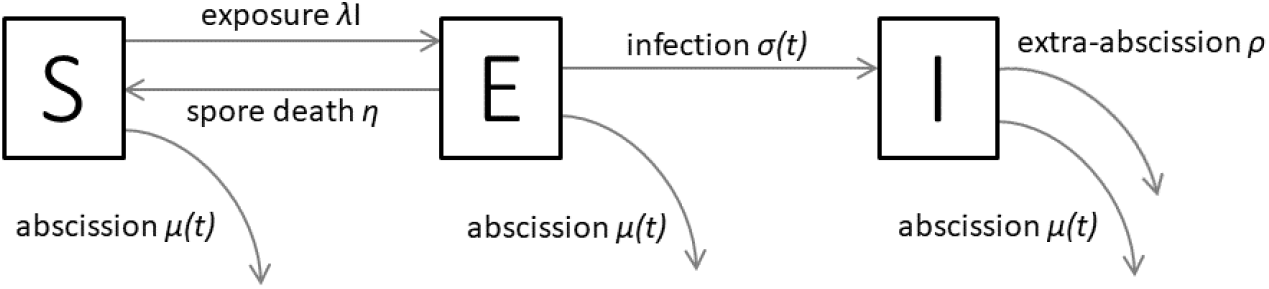
Schematic representation of the epidemiological model where, at each time *t*, fruits can be in either of the three states with respect to brown rot: susceptible (*S*), exposed (*E*) or infected (*I*). See text for details.

Because our analysis is extended to a two-year time span, the inter-annual fluctuations in fruit load can determine different fruit abscission rates between years, a factor that should not be neglected particularly in those years of high fruit production. In the current version of the system, we therefore explicitly modelled a fruit abscission rate. On the basis of considerations and analyses detailed in the Supplementary Information, the fruit abscission rate was modelled (and computed) as a function i) of fruit density, according to a power law, and ii) of fruit age, according to a parabolic relationship characterized by a null intercept (details in Supplementary Information).

### Model equations

According to the model outline and assumptions, the dynamics of the epidemics, from the time of pit hardening *t*_0_ to the harvest time *t_H_*, can be described with the following system of ordinary differential equations:

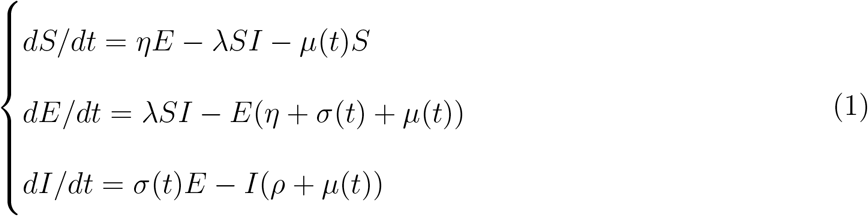

where *t* represents time, *η* the constant rate at which the spores deposited on the fruit surface die, λ the rate, per infectious unit, at which susceptible individuals are exposed to the pathogen, *μ*(*t*) is the abscission rate of fruits independent of their epidemiological state, *σ*(*t*) the rate at which a fruit exposed to the pathogen progresses to the infectious compartment, and *ρ* the extra-abscission rate of infectious fruits. All the model variables are referred to fruit density, *i.e*. number of fruits per 1 *m*^2^ of orchard surface. Given the regular geometry of plants in the considered orchard, the distance among trees was equal to 4 m, which corresponds to an average density of 625 plants per hectare [*ind ha*^−1^]. In practical terms, to compute the per tree fruit production it is simply a matter of multiplying the per area production [*kg m*^−2^] by 10^4^/625 [*m*^2^ *ind*^−1^]. Following Bevacqua et al. (2018), we assumed that

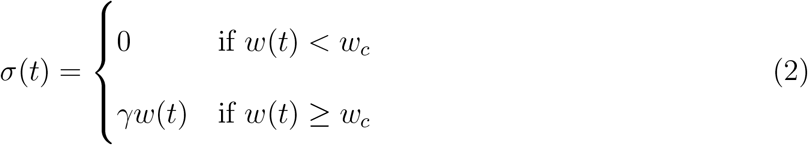

where *γ* is a constant progression rate toward abscission, *w*(*t*) is the fruit (fresh) weight at time *t*, and *w_C_* is the minimum fruit weight (threshold value) for infection being effective on it. The fruit weight *w*(*t*) is calculated as:

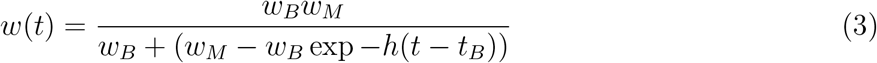

where *t_B_* is the time at fruit bloom, *w_B_* is the fruit weight at bloom, *w_M_* is the maximum fruit weight reached at the end of maturation and *h* can be interpreted as the conversion rate of resources into fruit weight (see Supplementary Information for details and Figure S8).

In principle, the abscission rate *μ* could be assumed to be dependent on either fruit age or fruit density (or both). After testing 29 competing models by means of the Akaike Information Criterion, the most promising model was the one were *μ* was computed as an (increasing) power law function of total fruit density *N* = *S* + *E* + *I* and a (concave down) quadratic function of the fruit age, computed as the difference between current time and time at bloom (i.e., *t* – *t_B_*, see Supplementary Information for details together with Table S1–S2, Figure S2 and S9). The equation expressing *μ* is therefore

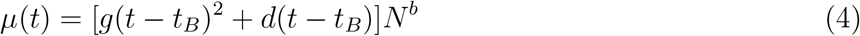

### Climate driven alternative models

Before performing a formal model selection on how climate dependent processes may affect brown rot dynamics in orchards, we explored the literature to screen which of the potentially many model parameters (each of which summarizing a process) were most credited to depend on weather (then climatic) conditions. After a careful scrutiny, we found that those were spore mortality rate *η*, exposition rate of susceptible fruit per infectious fruit λ and the progression rate *γ*, each of which was possibly varying with air temperature and/or precipitation events. When performing the formal model selection (see below) we systematically tested the alternative hypotheses of no dependence against dependence on each or both climatic variables.

The spore mortality rate *η* has been documented to increase with temperature (Xu and Robinson, 2000; Caubel et al., 2012). According to the Metabolic Theory of Ecology (Brown et al., 2004), the dependency on temperature of vital rates can be reasonably described by the Van’t Hoff-Arrhenius relation: exp (–*E_A_/kT*). We thus assumed two possible alternative expressions for *η*:

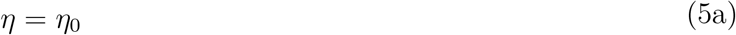

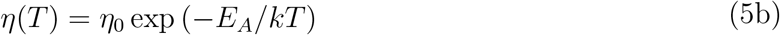

where *η*_0_ is the temperature independent part of *η*, *E_A_* (eV) is the activation energy of the considered process, *k* is Boltzmann’s constant (8.62 × 10^−5^*eVK*^−1^), and T is absolute temperature in K (Gillooly et al., 2001) (see Supplementary Information Table S3).

A key component of the exposure rate *λ* is the probability that a spore produced by an infectious fruit be dispersed nearby. Wind and precipitation are the two main drivers of spore dispersion (Fitt et al., 1989; Madden, 1997). As a function to be tested for dependence on climate variables, we therefore assumed a constant baseline exposure λ_0_ that could possibly be increased by a constant rate *l_P_*, that is positive only when precipitation occurs (*i.e*. *P* > 0). Said in plain words, the parameterization tested allowed pathogen exposure to increase during rain events due to the possible rain splash effect on the fungal spore dispersion. We disregarded precipitation intensity since spores dispersal occurs in the first minutes of precipitation events (Madden, 1997). In sum, we assumed two possible alternative expressions for *λ*:

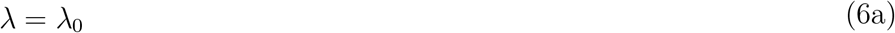

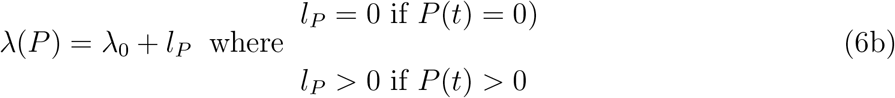

where *l_P_* represents the “splashing” component of the exposure rate.

It is likely that in plant-fungal diseases the infection rate (the part of it that was above called progression rate, *γ*, see eq. 2) can be activated by a precipitation event, probably because it increases fruit skin vulnerability and/or create suitable wetness conditions for fungal growth Caubel et al. (2015)). Other studies (Duthie, 1997) instead describe the progression rate as a unimodal (peaking) function of temperature. Following this rationale, we modeled the precipitation dependence (*α*(*P*)) and the temperature dependence (*β*(*T*)) of the progression rate as:

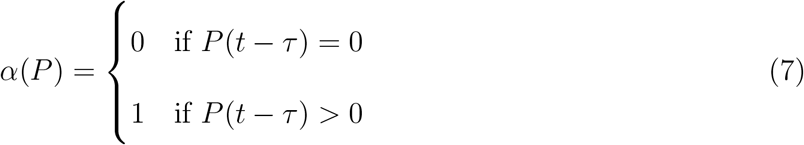

with *τ* corresponding to the temporal delay in days between the precipitation event and the effective increase in the progression rate *γ*, and

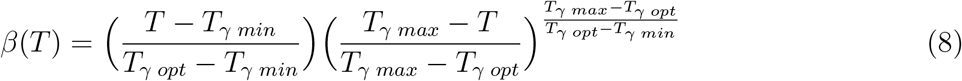

with *T_γ min_, T_γ opt_* and *T_γ max_* respectively representing the minimal, optimal and maximal average day temperature for fruit infection and *T* the daily average temperature (Yin et al., 1995). Accordingly, not only we investigated both hypotheses (*i.e*. dependence on temperature or precipitation), but also their combined effect. We therefore ran model selection among four possible expressions for *γ*:

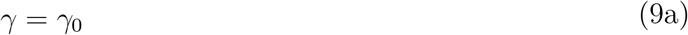

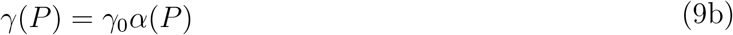

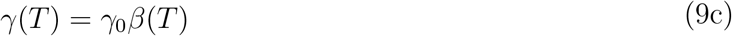

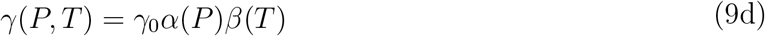

### Model calibration and selection

In order to minimize the number of parameters to be concurrently estimated from the available dataset, we kept fixed all parameters that had robust enough literature evidence to support that choice. For example, we set the critical fruit weight threshold for infection, *w_c_*, equal to 61 g (Gibert et al., 2007). Similarly, we fixed *T_γ min_* and *T_γ max_* to 6.5 °C and 23 °C, according to evidence by Xu et al. (2001) (see SI and Figure S10). For those (other) parameters that were instead estimated from the available dataset, we used a hierarchical scheme.

First, we calibrated the parameters that are related to three sub-models embedded in the overall modeling framework, namely: i) the natural fruit abscission (parameters *b, d* and *g*) by using observed abscission rates in absence of disease in 2014-2015 (number of observations=20), ii) fruit growth (parameters *w_B_*, *w_M_* and *h*) by using fruit weight measures across the 2014 and 2015 seasons (number of observations=2,737) and iii) temperature dependent spore mortality (*η*_0_ and *E_A_/k*) using observations from Xu et al. (2001) (number of observations=12) (see SI for details).

Second, by keeping fixed the parameters of the above mentioned three sub-models, we calibrated the remaining parameters, namely λ_0_, *γ*_0_, *ρ*, and the potentially climate-dependent components (*l_P_*, *T_γopt_* and *τ*), by running the full epidemiological model and using the 2014-2015 field observations of symptomless and infected fruit density (number of observations=26).

In the calibration procedures, we minimized the Sum of Square Errors *SSE*, with respect to the unknown parameters:

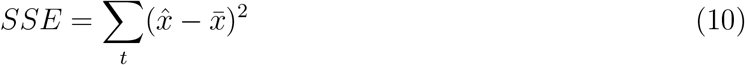

where 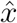 represents the estimated and 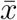 the observed data values.

When evaluating the model performance over the considered two growing seasons (2014-2015), we obtained the *SSE* summing the model performances of each year. We then assessed the empirical probability distributions of the estimated parameters by means of classical and moving block bootstrap, this last being recommended for time series (Kreiss and Lahiri, 2012). As for simulations performed with the complete epidemiological model, we numerically solved and simulated the infection dynamics from *t*_0_ to harvest time *t_H_*, by using the MATLAB’s solver for ordinary differential equations (`ode45`), which implements a Runge-Kutta method with a variable time step for efficient computation. We notice that the temporal scales of the steps are set by the units of measures of rates (in our case, time is measured in days). In accordance with field observations, and under the assumption that the exposed fruits at the initial phases (at *t*_0_) represent 1.8% of the entire fruit population (Emery et al., 2000), we set *S*(*t*_0_, 2014) = 8.94 (fruit m^−2^), *S*(*t*_0_, 2015) = 21.0 (fruit m^−2^), *E*(*t*_0_, 2014) = 0.16 (fruit m^−2^), *E*(*t*_0_, 2015)= 0.39 (fruit m^−2^), *I*(*t*_0_, 2014) = 0.03 (fruit m^−2^) and *I*(*t*_0_, 2015) =0.03 (fruit m^−2^).

To assess whether and how different models are better suited to reproduce the observed patterns, we ranked the performances of different candidate models according to the Akaike’s Information Criterion corrected for small samples (*AIC_c_*) (Burnham et al., 2002; Symonds and Moussalli, 2011).

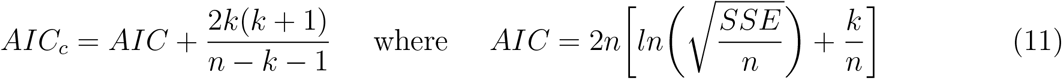

where *k* is the number of estimated parameters, *n* is the sample size and SSE is minimized in the calibration procedure (Symonds and Moussalli, 2011; Burnham et al., 2002). A complete description of competing models is reported in Table S4. According to Symonds and Moussalli (2011), we considered as equivalently good models whose *AIC_C_* scores differed by a *ΔAIC_c_* < 2 and, among them, we selected as best model the most parsimonious one or, in case of equal complexity, the one with lowest *AIC_C_*. We compared seven models for fruit abscission, two models for spore mortality and eight epidemiological models obtained by different combinations of eqs. 6–7. For the sake of simplicity and in line with the hierarchical calibration procedure described above, we performed subsequent Akaike model selections: *i*) we selected the two best sub-models of fruit abscission and of spore mortality, embedding them in the epidemiological model (*i.e*. eqs. 4 and 5); *ii*) we selected the best epidemiological model obtained from eight different combinations of eq. 6 (a–b) and 9 (a–d).

### Model sensitivity to climate variations and climate change impacts

We used the selected epidemiological model to estimate the consequences of brown rot disease on the annual yield in a virtual orchard subjected to different climatic conditions: *i*) as recorded in the past decade 2010-2019 in Avignon (Figures S3, S4); *ii*) as predicted for the 21st century (2030-2099), under different Representative Concentration Pathways (RCP) scenarios, in a 8 × 8 km^2^ cell comprehending Avignon (see SI for details regarding the climatic scenario generation and Figures S6, S7). Being the virtual orchard *i*) initialized every year with average conditions observed in 2014 and 2015 [*S*(*t*_0_) = 15.0, *E*(*t*_0_) = 0.28 and *I*(*t*_0_) = 0.03, all measured in fruit m^−2^], and *ii*) characterized by the same parametric setting, the yield magnitude only depends on different climate conditions. Climate can affect peach yield by *i*) impacting plant phenological times and fruit growth, thus determining the values of *t_B_*, *t*_0_ and *t_H_*, and *ii*) influencing the epidemiological processes of brown rot spreading. We estimated the timing of the peach growing season according to the phenological models calibrated for peach and presented in Vanalli et al. (2021). Being *t_H_* a cultivar-dependent parameter, we reassessed parameter *R_r_* (eq. 8 in Vanalli et al. (2021)) as equal to 1452 Growing Degree Days (GDD) for the cultivar Magic, used in the present work. Given blooming and harvest times for each growing season, we calculated the conversion rate *h* of the fruit logistic growth curve by inverting eq. 3, fixing the fruit weight at harvest to the average value at harvest in 2014 and 2015 (*w*(*t_H_*) = *w_H_* = 164*g*). In this way, the length of the fruit growing period, i.e. from blooming to harvest, determines how fast the fruit reaches harvest size. We assumed that the remaining parameters that describe the fruit logistic growth curve (*w_B_* and *w_M_*, see eq. 3) were constant for the Magic cultivar. As it was done in Bevacqua et al. (2018), we instead computed the yield as the fresh weight of symptomless, thus marketable, fruit at harvest time *t_H_* (*i.e*., *S*(*t_H_*) + *E*(*t_H_*)). To let the reader understand whether values are high or low compared to a reference value, we reported yield as the percentage of what it would have been in absence of disease in that same year. To evaluate future changes in the yield composition and amount under climate change, for each considered decade (i.e. 2030-2039, 2040-2049, 2050-2059, 2060-2069, 2070-2079, 2080-2089,2090-2099), we averaged the 10 annual yields and relative compositions, considering a null yield in those years when blooming failure occurred.

Eventually, to disentangle the effect of climate over phenological times from that over epidemic parameters, we performed a sensitivity analysis of the brown rot induced yield losses to variations in temperature and precipitations over a fixed time horizon (i.e. fixing *t*_0_ = 7*^th^* June and *t_H_* = 16*^th^* July whatever the climatic conditions). We varied temperature and precipitation occurrence up to +/- 20% compared to the average values recorded in the decade (2010-2019) (See SI for details).

## Results

### The climate-driven epidemiological model

As anticipated above, from the model selection described above, the fruit abscission rate resulted to depend on both fruit density and age via a power law and a parabolic function, respectively (details of the analyses are in the Supplementary Information, Table S2 and Figure S9), while the mortality for spores resulted to depend on temperature only (eq. 5b, see Table S3). In the second step of our hierarchical calibration method (see previous section), the optimal trade off between the calibration of full epidemiological model and the lowest complexity resulted in the selection of the model labelled as M3 (see Table S4 and Table S5), characterized by precipitation, but not temperature, dependence of the progression rate (*i.e*. eqs. 7 and 9b) and a temporal delay *τ* (eq. 7) of five days, between the occurrence of the precipitation event and the increase of the progression rate *γ* (see eq. 9b). A description of the selected model parameters and their units is summarized in Table 1.

**Table 1:** Model parameters, definitions, reference equations (Eq), estimated values, 90% confidence intervals (CI) and dimensions. The first seven parameters (from *w_B_* to *w_C_*) are specific to the host plant physiology (*Prunus persica*), the remaining six (from *γ*_0_ to *τ*) are specific to the considered disease (the brown rot).

| Parameter | Definition | Eq. | Value | 90% CI | Unit |
| --- | --- | --- | --- | --- | --- |
| $w_B$ | Fresh fruit weight at bloom time | 3 | 0.49 | [0.39;0.61] | g |
| $w_M$ | Maximum fresh fruit weight | 3 | 214 | [198;233] | g |
| $h$ | Conversion rate from resources to fruit mass | 3 | $5.6 \times 10^{-2}$ | $[5.3;5.9] \times 10^{-2}$ | d<sup>-1</sup> |
| $g$ | Fruit age square coefficient for fruit abscission | 4 | $-6.8 \times 10^{-7}$ | $[-81;-0.38] \times 10^{-7}$ | d<sup>-2</sup> |
| $d$ | Fruit age linear coefficient for fruit abscission | 4 | $9.1 \times 10^{-5}$ | $[2.5;86] \times 10^{-5}$ | d<sup>-1</sup> |
| $b$ | Fruit density exponent for fruit abscission | 4 | 0.49 | $[-1.9 \times 10^{-2}; 0.74]$ | - |
| $w_c$ | Fresh fruit weight threshold for cuticle cracking | 2 | 61 | [54;69]<sup>*</sup> | g |
| $\lambda_0$ | Exposition rate of susceptible fruit per infectious fruit | 6a | 0.15 | [0.13;0.20] | fruit<sup>-1</sup> m<sup>2</sup> d<sup>-1</sup> |
| $\rho$ | Extra abscission rate in infected fruit | 1 | $4.76 \times 10^{-2}$ | $[1.29;5.48] \times 10^{-2}$ | d<sup>-1</sup> |
| $\eta_0$ | Spore death normalization constant | 5b | $5.7 \times 10^{19}$ | $[1.1 \times 10^{16}; 1.4 \times 10^{26}]$ | d<sup>-1</sup> |
| $E_A$ | Activation energy | 5b | 1.2 | [0.9;1.7] | eV |
| $\gamma_0$ | Infection constant | 9b | $3.18 \times 10^{-3}$ | $[1.69;6.80] \times 10^{-3}$ | g<sup>-1</sup> d<sup>-1</sup> |
| $\tau$ | Delay between precipitation and infection rate increase | 7 | 5 | [4;6] | d |

\* estimated from Gibert et al. (2007)

Adherence of the selected climate driven model (M3) to field data is reported in Figure 2, while the performance of the original climate-independent model, with and without fruit abscission, is reported for comparison in the Supplementary Information (Figure S11).

**Figure 2:**
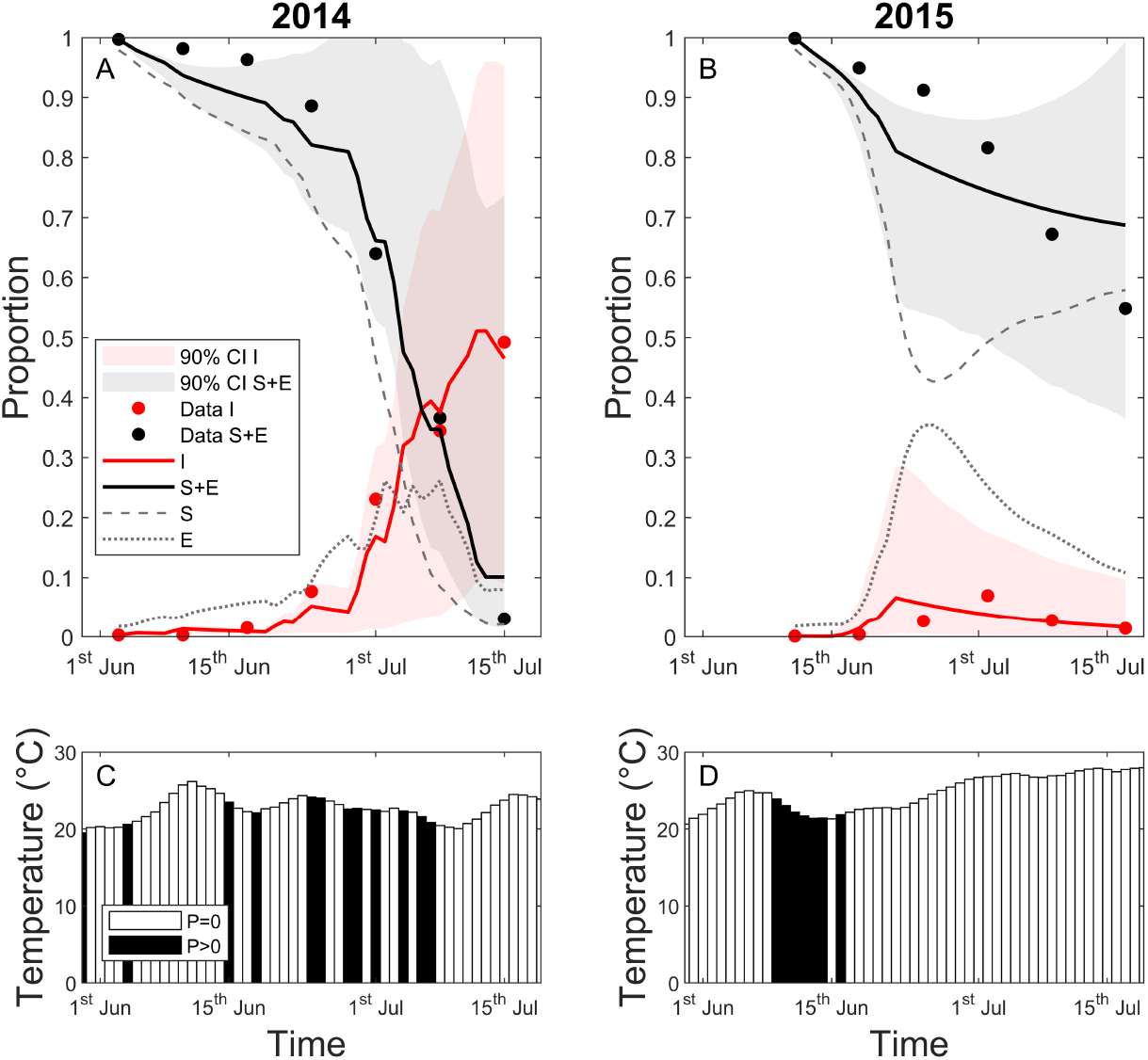
Temporal dynamics of the prevalence of susceptible (*S*), exposed (*E*), infected (*I*), and symptomless (*S* + *E*) fruit densities, simulated via the climate driven selected model of Table 1 (lines) and observed (circles), black for symptomless and red for infectious fruits) in 2014 (panel A) and 2015 (panel B). Shaded areas (same color coding) indicate the predicted 90% confidence bands for symptomless and infectious fruits, respectively. Observed daily average temperature, filtered by a weekly moving average, and precipitation occurrence (in black when *P* > 0) in 2014 (panel C) and 2015 (panel D).

Our best climate driven model fits well the epidemics trajectories not only in 2014 but also in 2015. In the Supplementary Information, (see Figure S11) we show that an equivalent but climate-independent model would not be able to fit observations of neither of the two years. Our climate-driven model therefore provides a mechanistic confirmation that the combined effect of absence of precipitation and high temperatures, in the four weeks preceding harvest, impeded the spreading of brown rot, in summer 2015. A quite continuous series of precipitation events occurred at the end of spring in 2015 determined indeed a first outbreak, but they were not maintained over the season of that year.

### Model sensitivity to climate variations and impacts of climate change

The consequences of temporal climatic variations, observed in the decade 2010-2019, over the considered peach-brown rot pathosystem are summarized in Figure 3. The estimated pit-hardening and harvesting dates, *t*_0_ and *t_H_*, significantly varied among years (Figure 3A) in response to different climatic conditions (see Figure S3). We estimated the earliest *t*_0_ corresponding to the 2^nd^ June in 2017 and the latest to the 1^st^ of July in 2013. Estimated *t_H_* varied between July 14^th^ in 2017 and August 6^th^ in 2013. Different climatic conditions are responsible also for the variation in the infection rate (Figure 3B) and in the spore death rate (Figure 3C), which affects the yearly brown rot spreading and eventually the yield composition. Our results indicate that climate conditions recorded in the last decade, with average daily temperature between 22.7-26.3 °C and 1-10 rainy days in the growing period [*t*_0_ – *t_H_*], could have determined disease-induced yield losses spanning from 4 to 81% in untreated orchards. Notably, year 2014, which was used to calibrate the original model of Bevacqua et al. (2018), is likely to have been the most impacted year by brown rot in the 2010-2019 period. In seven years out of ten of that decade, losses would have been below 6%. Although the number of rainy days is positively related with estimated losses due to brown rot, our simulations show that the timing of the precipitation events is crucial (see Figure S4). On the one hand, the dependence of the infection rate on the fruit weight determines a higher probability for a fruit to get infected at the end of the growing season, provided that there is a precipitation event. On the other hand, if favorable conditions for infection are met earlier in the growing season, there is more time for a polycyclic disease to spread and to cause higher yield losses before the time of harvest. Looking only at the number of rainy days, one would have expected higher losses in 2016 and 2019. However the fact that these rainy days were concentrated at the end of the growing season, in 2016, and that air temperatures were extremely high, in 2019, permitted to limit brown-rot induced losses. The small differences in potential yields, i.e. in case of disease absence, from 2.00 to 2.12 kg/m^2^ is due to the fact that fruit cumulative abscission rate increases with growing season duration.

**Figure 3:**
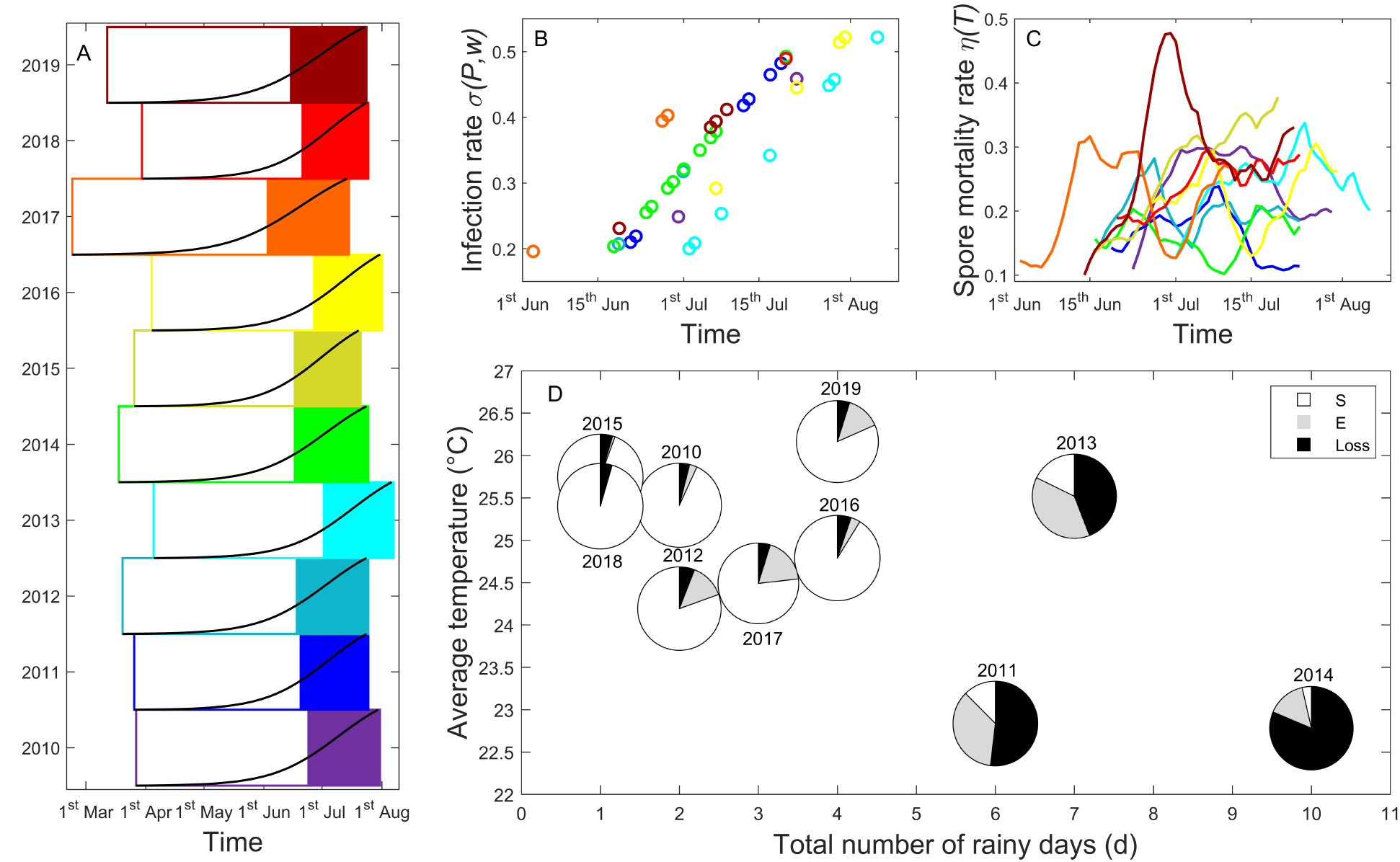
(Panel A) Estimated phenological times for peach (Magic cultivar) over the decade 2010-2019, with different years characterized by different colors. For each year, the empty fraction of the rectangle represents the time from blooming *t_B_* to pit hardening *t*_0_ and the filled one the time from *t*_0_ to harvest time *tH*. The black line inside each rectangle represents the fruit growth curve. (Panel B) Predicted infection rate *σ*(*P, w*) over the considered growing season (x-axis). Infection rate is activated 5 days after a precipitation event, *P* > 0, and increases with fruit fresh weight w. Each color identifies a given year, as coded in panel A. (Panel C) Predicted spore mortality rate *η*(*T*), over the considered growing season (x-axis), increasing with temperature. Each color identifies a given year, as coded in panel A. (Panel D) Estimated yearly yields, from 2010 to 2019, in relation to the total number of rainy days (pie center abscissa) and the average temperature (pie center ordinate), recorded during the peach growing season (i.e. from *t*_0_ to *t_H_*). White and gray pie slices indicate the fraction of Susceptible (S) and Exposed (E) fruits at harvest, respectively, and their sum represents the estimated yield in presence of disease. The black pie slices are a measure of the disease-related losses. The size of each pie is proportional to the estimated annual yield in absence of disease, which ranges from a minimum of 2.00 kg/m^2^ to a maximum of 2.12 kg/m^2^, respectively in 2017 and 2015.

Once the mechanistic model with climate-relevant parameters has been properly calibrated and show credible results for a decade, we can use it to investigate what could be the effects of climate change over a longer time horizon. The consequences of predicted climate changes in the current century (see SI Figure S6, S7) for peach phenology and yield, both in absence and presence of brown rot, are reported in Figure 4. Our results indicate that the years of absence of yield, due to unsuitable conditions for peach tree cultivation, will increase over the century and will be more abundant in the 8.5 RCP scenario (Figure 4A). This is mainly due to the fact that too warm winters will impair blooming not permitting the fulfillment of chilling requirements (see Vanalli et al. (2021) for details). There is no clear trend on the impact of brown-rot related losses over the century (black portion of the pies in Fig. 4), with the exception of a smaller percentage of losses in the last decade in the RCP 8.5 scenario. That decrease is probably due to significantly warmer and drier conditions. However, rather than the changes in fractions of yield losses, the most important result that emerge from the analysis is the reduction of the potential yield (areas of pies) over the decades. This is particularly true in the RCP 8.5 scenarios and it is an important warning that, at least for the presented cultivar and in the considered Mediterranean region of southern France, climate changes will be likely to impact peach production more by making conditions unsuitable for peach cultivation than by increasing the risk of brown rot. This is due to the fact the number of rainy days, in the predicted growing seasons, which affects brown rot incidence, is not predicted to significantly change. However, on the other hand, the increase in temperatures over winter and spring seasons is likely to have important impacts on the plant phenology possibly impairing blooming. The negative effect of warmer and drier conditions over the epidemic spreading is shown in detail in Figure S12. In the reference scenario, corresponding to the average conditions observed in the decade 2010-2019, the expected losses are 21% with respect to a potential yield, in case of disease absence. For RCP 4.5, such a value would range from 20% up to 44%, while for RCP 8.5 from 9% to 51% (Fig. 4C).

**Figure 4:**
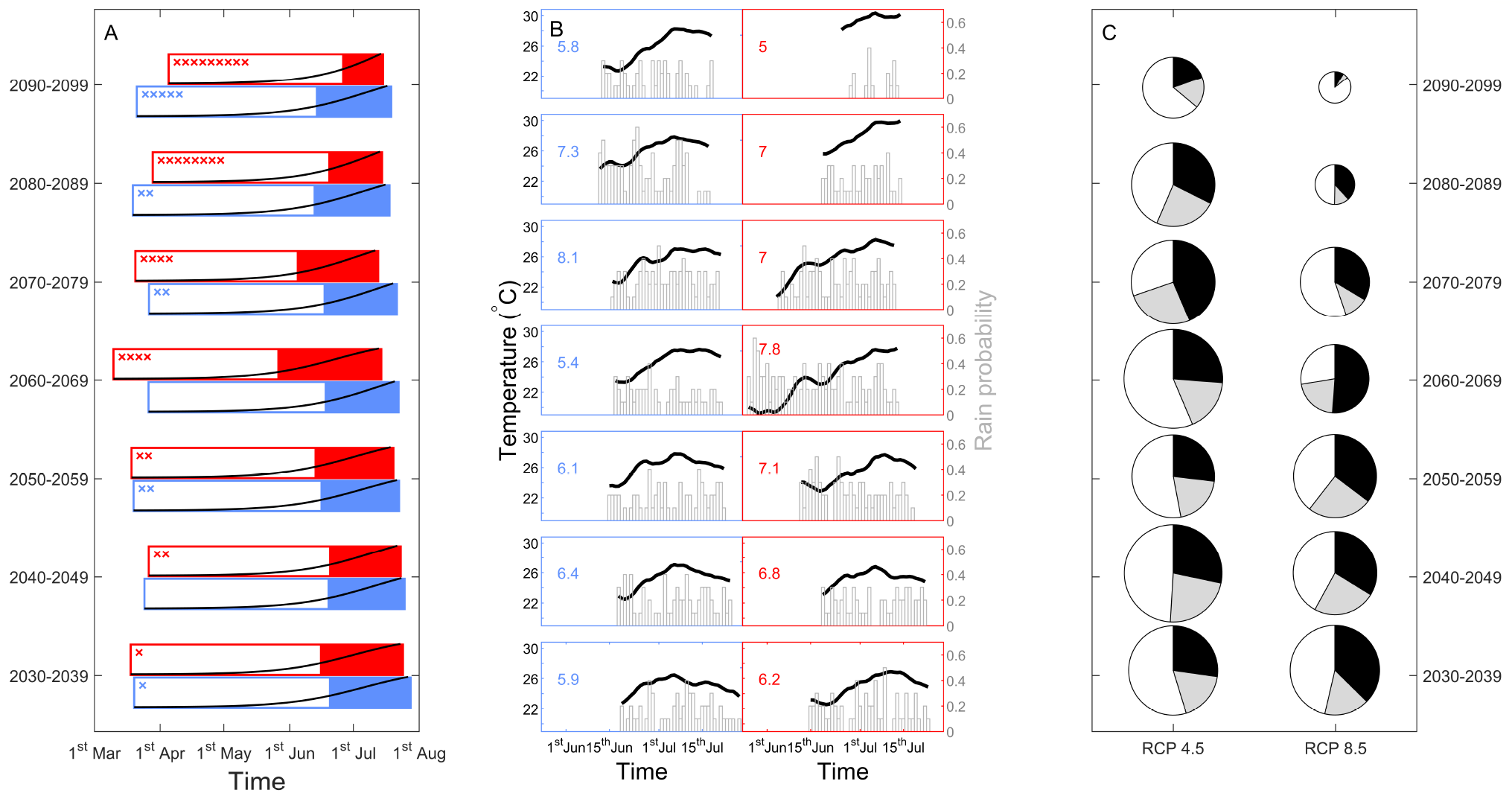
(Panel A) Average phenological times of the peach growing season, under RCP 4.5 (in blue) and RCP 8.5 (in red) scenarios, for different decades across the 21st century (left axis). As in Figure 3, the empty fraction of each rectangle represents the average time from blooming *t_B_* to pit hardening *t*_0_, while the filled one indicates the average time from *t*_0_ to harvesting time *t_H_*. The black lines represent the average fruit growth curves. The number of years within each decade when peach blooming failed to occur because of climate-dependent reasons (see Vanalli et al. 2021) is reported by the “x” symbols. (Panel B) Daily average temperature (in black, left axis) and rain probability (in gray, right axis) in the growing period from *t*_0_ to *t_H_* for each decade under RCP 4.5 (blue rectangles) and RCP 8.5 (red rectangles) scenarios. The average annual number of rainy days in the considered period is reported. (Panel C) Average annual yield in different decades across the 21^st^ century, under RCP 4.5 and RCP 8.5 scenarios. White and gray pie slices represent the estimated yield proportion of Susceptible (S) and Exposed (E), while the black slices are the brown rot-related losses, compared to the yield in absence of disease. The size of each pie is proportional to the average annual yield in absence of disease in a specific decade, from a minimum of 0.22 kg/(m^2^ year) to a maximum of 2.2 kg/(m^2^ year).

## Discussion

Our results indicate that including a temperature dependent spore mortality and a precipitation dependent infection rate in a SIR-type model, properly adapted to study brown rot in peach orchards, allows to reproduce very different epidemic trajectories observed in two subsequent years (Fig. 2). The model studied here is extended after Bevacqua et al. (2018) and analyzed by accounting for different meteorological conditions. It successfully permitted us to simulate the changes in phenological times of the host, the severity of epidemic outbreaks and consequent brown rot-related yield losses over a past 10-year period (2010-2019, see Fig. 3). Also, the model permitted us to forecast the consequences of climate change, in the study area, for the 21st century under the two RCPs scenarios 4.5 and 8.5 (Figure 4).

Our results could be contrasted against field data, but only for years 2014 and 2015, when they proved to well fit what happened. A sound validation of our decennial analysis would require in fact disease observations over a longer temporal scale that are, to date, still unavailable. Therefore, we agree with Bebber et al. (2016) when saying that “investment in monitoring, storage and accessibility of plant disease observation data are needed to match the quality of the climate data now available”. Their consideration is also valid to let researchers evaluate reliability of simulated scenarios. Nowadays, efforts to provide open databases to serve scientific research are increasing.

This is even more evident with regard to epidemiology, especially when it directly involves humans. Just think of the many articles on COVID-19 based on the open access database provided for instance by the Johns Hopkins University (https://coronavirus.jhu.edu/). Although this trend is more evident for human diseases, something similar is occurring also for crop diseases. For instance, the Plants section of the ISID-ProMED database https://promedmail.org/ has been recently used by Romero et al. (2022) to evaluate the relative importance of different climatic drivers for fungal pathogen outbreaks (but see Bebber (2022)). However, for brown rot in France, we could not find either qualitative or quantitative indicators of annual losses in different years and locations. Available data are gathered by different institutions and with different protocols and different aims. In order to possibly use them, an intensive work of research, data grasping, data analysis and homogenization must be done. Such a work is out of the aim of the present study. While such work would certainly give some useful information, it would require a huge amount of effort without certainty about its usefulness. Otherwise, the use of common protocols, devised by the scientific community in collaboration with the various institutes and/or producers, would be of enormous help to research and understanding of how agro-ecological systems work.

In general terms, our results indicate that the expected future warmer temperatures and fewer rainy days, with respect to the historical trends of a northern Mediterranean region, would result in lower percent losses due to brown rot but must be coupled with a reduced yield due to climate change. Our epidemiological findings are in contrast with those of Desaint et al. (2021), who argued that temperature increase will globally augment the number and severity of crop epidemics, regardless of the plant and pathogen species. Such misalignment of previsions is less effective than it might appear, because the above mentioned findings are mainly due to predicted trends in the temperate regions of Europe, where increased temperatures will match the optimal requirement of the considered pathogens (Chaloner et al., 2021). In accordance with our results, Bregaglio et al. (2013) found that climate warming might determine a lower disease incidence for those regions where temperature will exceed optimal ones with respect to the pathogen and Tresson et al. (2020) identified a reduced blossom blight risk for those cultivars that will be exposed to future drier and warmer conditions. Our findings also show that the timing of occurrence of favorable environmental conditions for brown rot spreading, i.e. within season frequent rain occurrence and low temperatures, influence the infection success and the consequent yield (Fig. 3 and Fig. S4). On the one hand, the interplay between the host growth and the pathogen dispersal suggests that infection tends to be higher at the end of the growing season: the infection rate is proportional to the fruit size and negligible for fruit which has not overcome a critical size threshold, with fruit size increasing along the season following a sigmoidal trajectory (see Figure S8). On the other hand, if the infection is triggered by a precipitation event at the beginning of the season, as soon as the fruit crosses the weight threshold that allows infection, the epidemic has more time to spread before harvest. Overall, in case of rain occurrence, we expect higher disease severity when *i*) there are more rainy days within the growing season (frequency is more important than intensity); *ii*) rainy days are dispersed across the growing season; *iii*) concentrated rain events (i.e. consequent rainy days) occur early in the season, but following the day when fruits are louder than the weight threshold for infection (Fig. 3 and Fig. S3)..

Mechanistic models of weather-dependent disease risks in crops have been already introduced in the literature and used to develop disease forecasting tools. However, in most of cases (but see (Chaloner et al., 2019)) authors use infection models whose structure and parameters are not inferred from population data, neglecting the state of the host and focusing on the evaluation of the weather effects on the pathogen vital requirements (see e.g. Magarey et al. (2005); Bregaglio et al. (2012); Bebber (2019); Chaloner et al. (2019)). In the present work, we embedded approaches developed and used in infection models within a compartmental epidemiological SIR-type model, which allows us to consider also the temporal dynamics of the host status. In our model, the key variable determining the host status is the size of the fruit which increases over the season in a sigmoidal way. Incorporating epidemiology into a plant/host growth model, explicitly considering the effects of climate and agricultural practices, to assess crop yield/losses is a pivotal challenge for plant science (Cunniffe et al., 2015a; Donatelli et al., 2017; Zaffaroni et al., 2020, 2021). The study of the response of epidemiological systems, described via compartmental SIR or SEI-like models, to a varying (in time) environment has been already done for humans (Casagrandi et al., 2006; Rinaldo et al., 2012), animal (Bolzoni et al., 2008; Samuel et al., 2011) and plant (Truscott and Gilligan, 2003) diseases. Up to our knowledge, this work represents a first attempt to describe a crop epidemic system via a SIR-type formalism where the effects of climatic conditions are accounted for over both the host and the pathogen dynamics. Despite the limitations of our work, we believe it could represent a basis for forthcoming research aimed at using reliable scenarios of climate changes as drivers to project and analyze future crop production within a well established conceptual and theoretical framework for the study of epidemics.

In the present work, we fed our mechanistic model with two climate-related inputs, that are daily average air temperature and a Boolean variable (presence/absence) of daily precipitation occurrence. We chose these two weather variables because their combined effect had already been related to the *Monilinia* incidence risk (Tresson et al., 2020) and showed to promisingly grasp the core dynamics of the system without the need of using further, more refined climate variables. Also, making reference to variables that can be easily measured, our model results to be easily extendable and applicable to contexts where the availability of climate measurements is scarce. We are well aware that *i*) the microclimate conditions actually experienced by microorganisms determining plant diseases differ from those measured by weather stations and even differ within the plant canopy, and *ii*) many responses of biological rates to climate are non-linear, and the mean response is not the same as the response to the mean climate (see Bütikofer et al. (2020) for a recent review on this subject). Some studies on fungal infections also relate infection risk to moisture or wetness duration (see e.g. Bebber et al. (2016); Bebber (2019)). However, these more refined variables are not commonly measured and their estimate, even when available, is associated to high uncertainty (Bebber, 2015). Uncertainty also exists for the estimates of precipitation in future weather scenarios (Bregaglio et al., 2013). Yet, it is a variable intrinsically accounted for and whose spatial variability is much lower than that of moisture and wetness which can significantly vary even within a plant canopy. Of course, future improvement could also derive microclimate conditions in different parts of a plant as a function of environmental conditions and plant development (e.g. size and phenological stage). In particular, the inclusion of the plant and fruit growth responses to climatic stresses, such as heat and drought events, could provide more informative conclusions (Génard and Huguet, 1996; Desaint et al., 2021). Yet,at the level of a first approximation where the plant architecture is not explicitly considered here, our simplified modelling still hold to satisfactorily capture the effects of climate on fungal disease dynamics. Moreover, the proposed modeling formalism is suited to make the model parameters depending on additional (and in case even different) climatic variables.

Our work intends to simulate how climate variables affect plant diseases spread as they emerge from the dynamical interaction of hosts and pathogens. We assume that a primary inoculum of the pathogen is present in the system and we wonder if, and to what extent, the climate conditions will permit the disease to spread. We are aware that the fungal density of the primary inoculum at the beginning of the fruit growth season can itself depend on inter-seasonal climatic conditions, and thus it can produce different yield-related disease outcomes. However, estimating the intensity of a primary inoculum is technically very challenging and setting its dependence on climate is beyond the aim of the present work. Other mathematical tools to assess possible geographical shifts in pathogen distribution due to climate changes are known as Species Distribution Models (SDMs). These models aim to predict potential species distributions by matching species’ environmental (primarily climatic) preferences with conditions in physical space (see e.g. Bebber (2015); Bebber et al. (2014) for the spread of crop pest and pathogens and Vanalli et al. (2021) for the predicted geographical shift of commercial fruit trees). In perspective, one could use SDMs models to derive initial conditions of our epidemic model. In other words, one could imagine that the initial abundance of infected and exposed fruits depends on the outcomes of SDMs models.

Foreseeing the consequences of climate change scenarios over the dynamics of crop fungal pathogens is far from being an easy task. Our results show the existence of a clear relationship between disease severity and average climatic conditions during the growth season (Fig. 3). Future predictions are certainly complicated by the inner uncertainty related climate forecasts and to the pathogen adaptation to new climatic conditions (De Crecy et al., 2009; Boixel et al., 2022). Despite the inherent limitations of any modeling exercise, we are convinced that the proposed modeling framework may represent a significant step towards the dynamic representation of epidemiological processes in crop systems that will certainly have to deal with climate changes.

## Acknowledgments

Part of this work has been performed when CV was graduating in Environmental Engineering at Politecnico di Milano and making her master thesis in collaboration with the INRAE within ECOV-ERGER project https://ecophytopic.fr/recherche-innovation/proteger/projet-ecoverger. This action is led by the Ministry for Agriculture and Food and the Ministry for an Ecological and Solidary Transition, with the financial support of the French Biodiversity Agency on “Resistance and Pesticides” research call, with the fees for diffuse pollution coming from the Ecophyto plan. The work is also part of the INRA Metaprogramme ACCAF (project 429 CLIF), that the authors acknowledge. We thank Signoret V., Leon I., and Tran A. for field work, Quilot-Turion B. and Launay M. for discussion. We are grateful to the IE UMR 1114 team for taking care of the experimental orchard. Howerton E. is also thanked for her help in proofreading this paper.

## Author contributions

DB conceived the research and conducted fieldwork. DB, CV and MG analysed data and designed the model (calibrated and simulated by CV). DB, CV, RC and MG interpreted the results and wrote the manuscript.

## Supplementary Information

### Available data

#### Fruit growth data

From the beginning of May until mid-July in 2014 and 2015, fruit fresh weigh was measured in the experimental orchard (Avignon, southern France). Fruit were sampled from the trees on the borders of the orchard that were excluded from the observation of the epidemiological dynamic. Over the course of the growing season of 2014 and 2015, the fresh weight of 713 and 2,024 total fruits was sampled, respectively (Figure S1).

**Figure S1:**
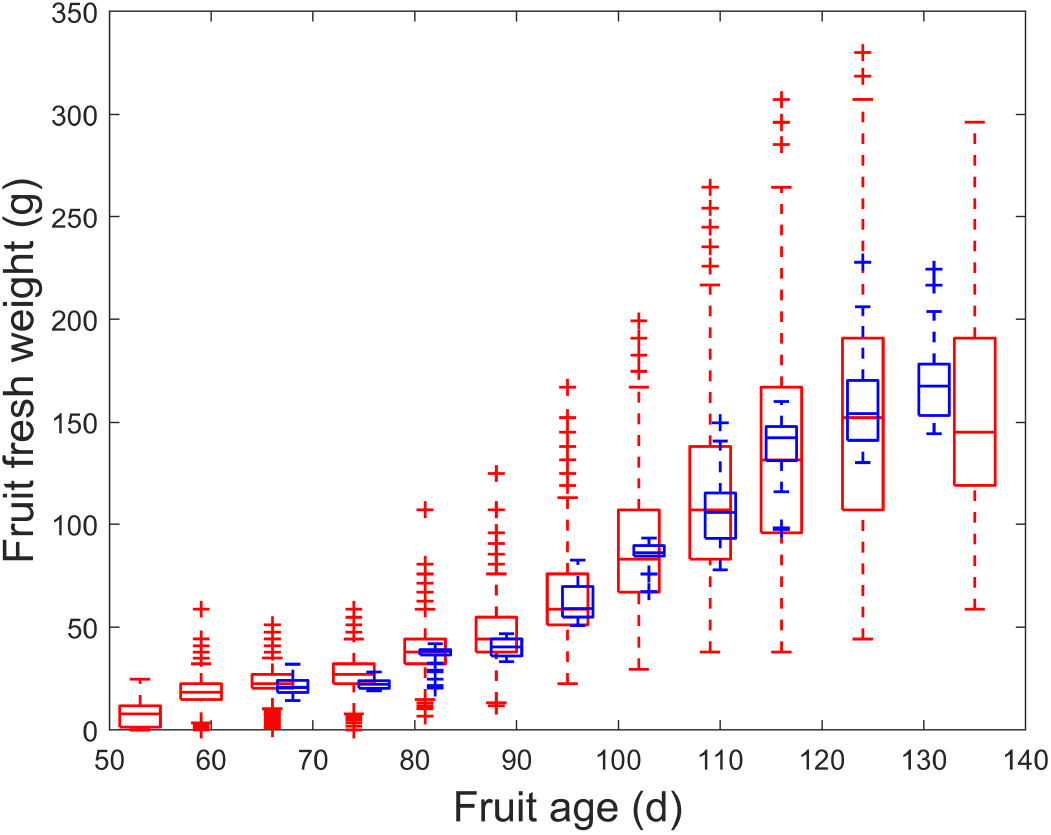
Observed fruit fresh weight (g) vs. fruit age in days after bloom (d). Blue and red boxplots represent observed data in 2014 and 2015, respectively.

#### Fruit abscission data

The fruit abscission rate can be estimated from the weekly observations of the variation of fruit densities, by building a sort of “life table”thus following procedure similar to that used for the calculation of the mortality rate in a population of individuals. For this purpose, we use dataset of both 2014 and 2015 fruit abundance, considering only data where the number of infectious fruits is less than 5% of the entire fruit sample, since, in presence of the disease, it would not have been possible to disentangle the natural falling rate (abscission *μ*) from the one induced by the infection (our parameter *ρ*). In addition, the orchard has been subjected to manual thinning on 17^th^ May (i.e. 137 Day Of the Year (DOY)) in 2014, and on 2^nd^ June (i.e. 153 DOY) in 2015, so the abscission rate cannot be estimated in the period 133-140 DOY for 2014 and 148-155 DOY for 2015. To calculate the mortality rates, we first applied data smoothing by averaging the logarithm of fruit density by a moving window that includes the previous (at *t*-1) and the following (at *t*+1) observations and then we estimated the abscission rate as

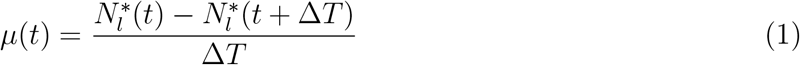

where *t* represents DOI, 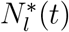 is the natural logarithm of fruit density at time *t* and *μ*(*t*) is the resulting abscission rate(d^−1^). We report the estimated abscission rates for 2014 and 2015 in Table S1.

**Table S1:** Dynamics of fruit densities (sort of “life table”) used to estimate the abscission rates *μ*(*t*) (last column), where *t* represents the Day Of the Year (DOY), *N*(*t*) represents the fruit density at day *t*, *N_l_*(*t*) its natural logarithm, 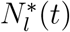 the smoothed logarithm of fruit density at day *t*.

| 2014 |  |  |  |  |
| --- | --- | --- | --- | --- |
| t<br>(DOY) | N(t)<br>(fruit m<sup>-2</sup>) | N<sub>l</sub>(t)<br>(-) | N<sub>l</sub><sup>*</sup>(t)<br>(-) | μ(t)<br>(d<sup>-1</sup>) |
| 100 | 21.6 | 3.08 | 3.08 | 5.3×10<sup>-3</sup> |
| 105 | 21.1 | 3.04 | 3.04 | 7.2×10<sup>-3</sup> |
| 112 | 20.5 | 3.02 | 3.00 | 1.3×10<sup>-2</sup> |
| 119 | 18.6 | 2.92 | 2.91 | 1.5×10<sup>-2</sup> |
| 126 | 16.2 | 2.78 | 2.80 | 1.5×10<sup>-2</sup> |
| 133 | 14.9 | 2.70 | 2.70 | - |
| 140 | 11.3 | 2.42 | 2.42 | 1.7×10<sup>-2</sup> |
| 147 | 9.8 | 2.28 | 2.30 | 1.1×10<sup>-2</sup> |
| 154 | 9.1 | 2.21 | 2.23 | 4.2×10<sup>-3</sup> |
| 161 | 9.0 | 2.20 | 2.20 | 1.3×10<sup>-3</sup> |
| 168 | 8.9 | 2.19 | 2.19 | - |

Table S1: Dynamics of fruit densities (sort of “life table”) used to estimate the abscission rates $\mu(t)$ (last column), where $t$ represents the Day Of the Year (DOY), $N(t)$ represents the fruit density at day $t$ , $N_l(t)$ its natural logarithm, $N_l^*(t)$ the smoothed logarithm of fruit ...
| 2015 |  |  |  |  |
| --- | --- | --- | --- | --- |
| t<br>(DOY) | N(t)<br>(fruit m<sup>-2</sup>) | N<sub>l</sub>(t)<br>(-) | N<sub>l</sub><sup>*</sup>(t)<br>(-) | μ(t)<br>(d<sup>-1</sup>) |
| 114 | 79.6 | 4.38 | 4.38 | 1.7×10<sup>-2</sup> |
| 120 | 74.7 | 4.31 | 4.28 | 2.3×10<sup>-2</sup> |
| 127 | 63.0 | 4.14 | 4.12 | 2.8×10<sup>-2</sup> |
| 134 | 48.8 | 3.89 | 3.92 | 2.6×10<sup>-2</sup> |
| 141 | 41.7 | 3.73 | 3.74 | 1.9×10<sup>-2</sup> |
| 148 | 36.9 | 3.61 | 3.61 | - |
| 155 | 23.6 | 3.16 | 3.16 | 1.1×10<sup>-2</sup> |
| 162 | 21.5 | 3.07 | 3.08 | 9.1×10<sup>-3</sup> |
| 169 | 20.5 | 3.02 | 3.02 | - |

The estimated fruit abscission rate *μ*(*t*) (see Table S1), depend on fruit density *N* (in fruit m^−2^) and on fruit age *A* (days after bloom) as shown in Figure S2.

**Figure S2:**
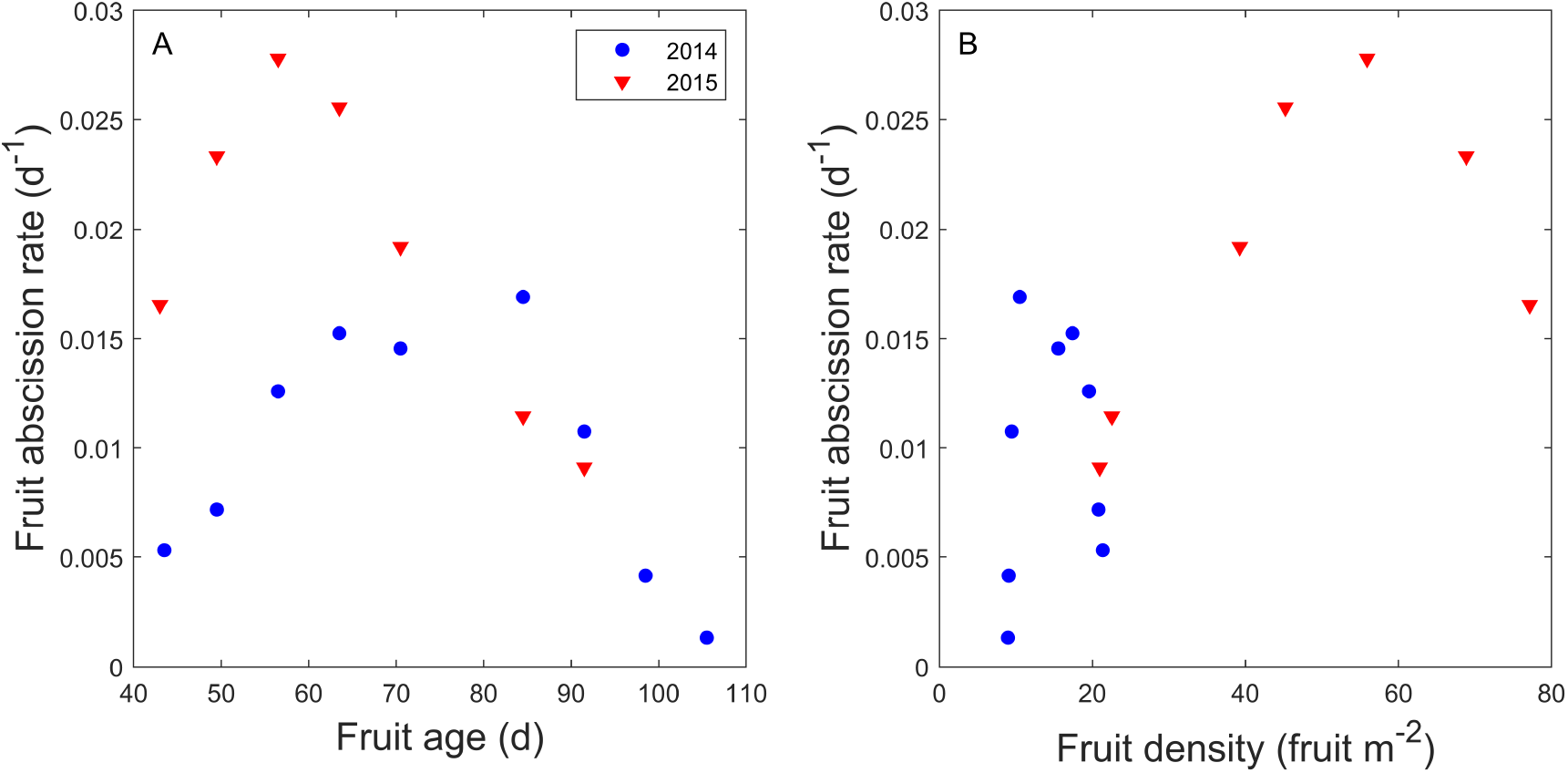
Fruit abscission rate dependency on (A) fruit age (days after bloom) and on (B) fruit density (fruit m^−2^)

#### Spore viability data

We used published data from Xu et al. [7] on spore viability at different temperatures, to estimate the mortality rate of the fungal spores. Xu et al. [7] observed spore viability at two different temperature conditions, 10 °C and 20 °C.

#### Climate historical data and generated climatic scenarios

For the time period 2010-2019, we obtained daily average temperature (°C) and daily precipitation intensity (mm) from the meteorological station of INRAE Saint-Paul (43.92°N; 4.88°E), see Figure S3.

**Figure S3:**
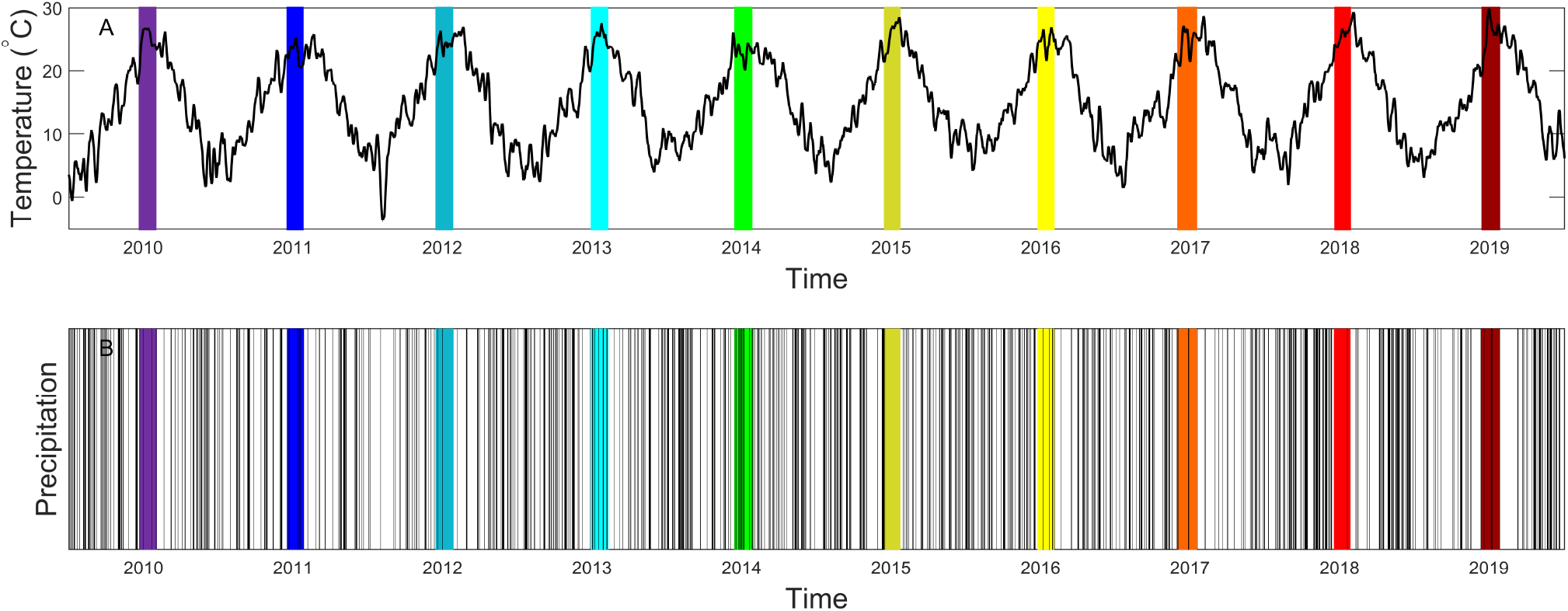
(Panel A) Daily mean temperature (°C), averaged with a moving window of ten days, and (Panel B) daily precipitation occurrence, from 1^st^ January 2010 to 31^st^ December 2019. For each year the time from pit hardening *t*_0_ to harvest *t_H_* is colored, with each color representative of a given year following color code of Figure 3 of the main text.

Climatic data during peach growing periods, estimated through phenological models, for each of the ten years are reported in in Figure S4.

**Figure S4:**
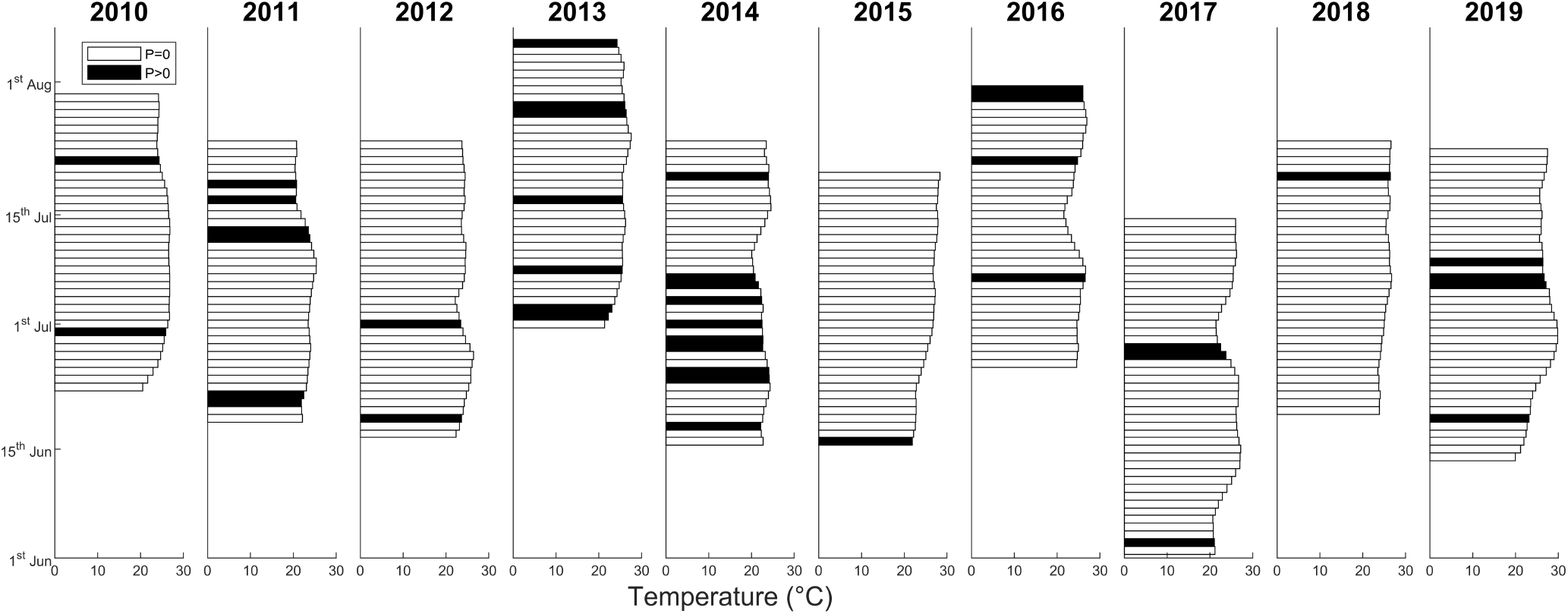
Daily mean temperature (°C), averaged with a moving window of ten days, and daily precipitation occurrence (*P* > 0) in the growing period from pit hardening *t*_0_ to harvest time *t_H_*.

To generate temperature scenarios to test model sensitivity to variations in temperature and precipitation, we increased/decreased the average daily temperature, observed in the 2010-2019 period, within the range of -/+ 20% (see Figure S5A). To reproduce plausible patterns of pre-cipitation occurrence, we derived daily probabilities of not having a precipitation (i.e. having a dry day *D*), followed by that of having a day with precipitation (i.e. having a wet day *W*), *P_DW_*(*t*), together with the probability of having two consequent precipitation days *P_WW_*(*t*), being *P_DD_*(*t*) = 1 – *P_DW_*(*t*) and *P_WD_*(*t*) = 1 – *P_WW_*(*t*), evaluating the frequency of these events per each day of the peach growing period over the ten years (2010-2019) of precipitation data [6].

For each value *v* of precipitation variation within the interval -/+ 20%, we generated 1,000 scenarios using a first-order Markov chain. More precisely, i) we extracted for each day *t* a random number *u* from a uniform distribution (*u* ∈[0;1]); ii) if the previous day (*t* – 1) was dry, then day *t* is simulated to be wet if *u* ≤ *P_DW_*(*t*) + *v* × *P_DW_*(*t*), and otherwise it is also dry; iii) if the previous day was wet, then the current day is simulated to be wet if *u* ≤ *P_WW_*(*t*) + *v* × *P_WW_*(*t*) and is dry otherwise; iv) initial conditions regarding the first day of the growing period (7^th^ June) are drawn with the probability of having a wet day *P_W_*(7^th^ June)+*v* × *P_W_*(7^th^ June), calculated from the ten years data (see Wilks and Wilby [6] for further details). The daily probabilities of having a dry day followed by a wet one *P_DW_*(*t*), together with the daily probabilities of having a wet day followed by a wet one *P_WW_*(*t*) and their +/- 20% changes are reported in Figure S5B. We evaluated all the combinations between temperature scenarios and precipitation occurrence scenarios, noting that one temperature scenario corresponds to 1,000 stochastic scenarios of rain occurrence.

**Figure S5:**
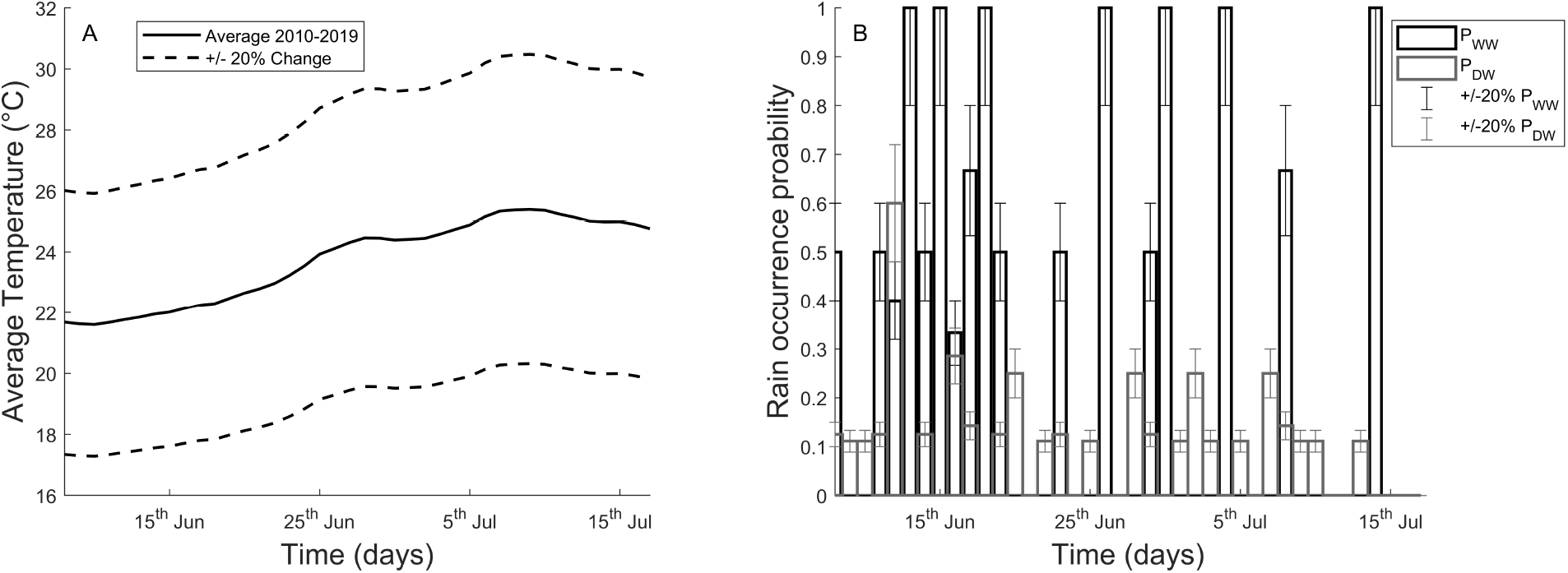
(A) Daily average temperature (°C) in terms of daily mean (solid) and a variation +/- 20 % (dashed). (B) Daily probabilities of having a dry day followed by a wet one (*P_DW_*(*t*), in blue), together with the daily probabilities of having a wet day followed by a wet one (*P_WW_*(*t*), in red). Even for the precipitation, the recorded daily average are represented together with the +/- 20% changes of these variables.

#### Future climatic scenarios: Representative Concentration Pathways 4.5 and 8.5

Future daily average temperature and precipitation occurrence time series across the 21^st^ were generated by the project Drias, developed by Meteo-France, with a spatial resolution of 8 × 8 km^2^ (Figure S6, S7). We used future climatic projections of two Representative Concentration Pathways, an intermidiate scenario, RCP 4.5, and a business as usual scenario, RCP 8.5 (IPCC, 2013). The two RCPs refer to different future green house gas emission trajectories, corresponding to a value of the radiative forcing at the end of the century of 4.5 W/m^−2^ and 8.5 W/m^−2^, respectively. For daily average temperatures, we performed a statistical downscaling of future projections in the Avignon site, applying the delta addition method based on the historical average daily temperatures observed from Avignon meteorological station, from 2010 to 2019.

**Figure S6:**
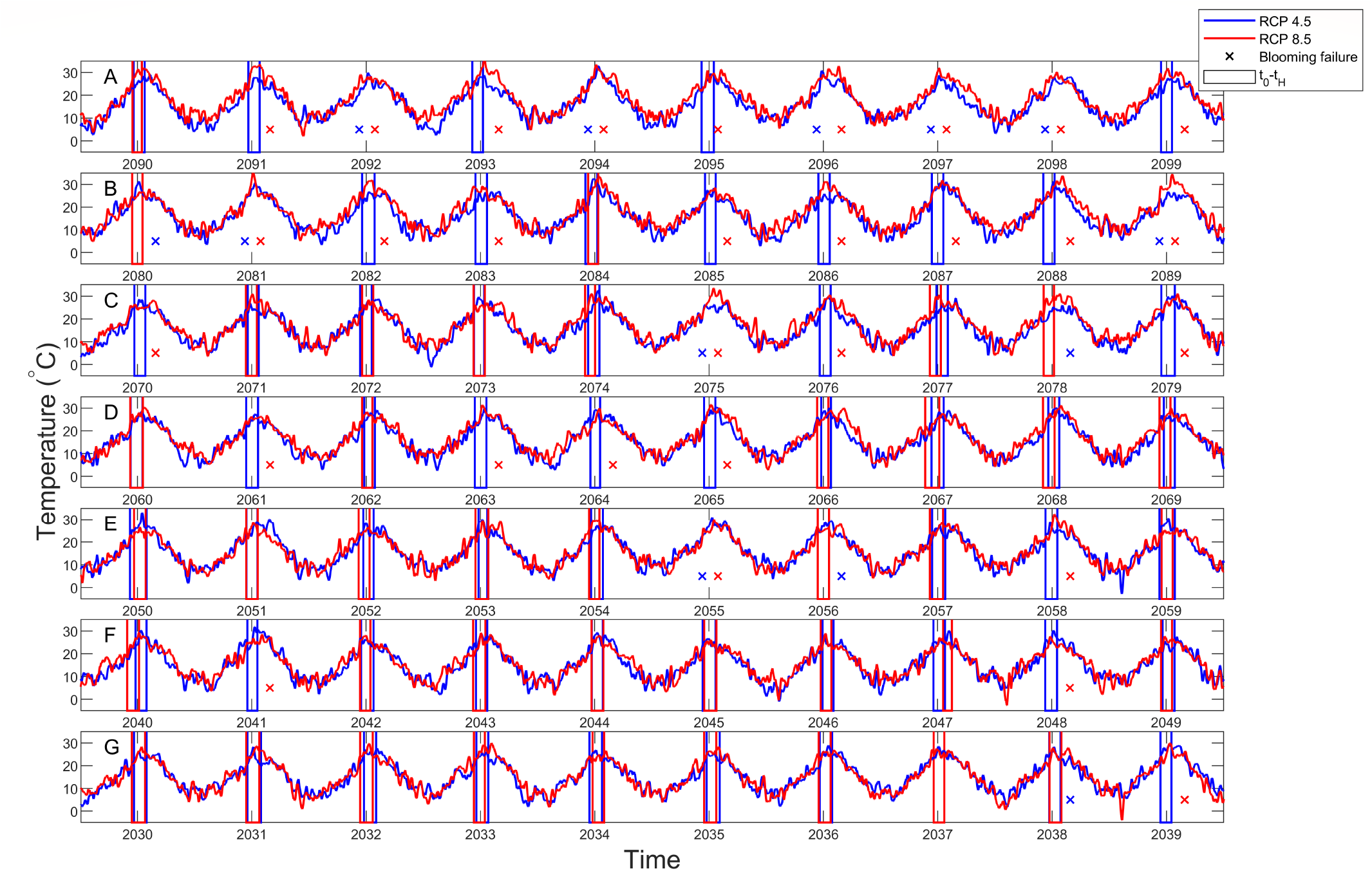
Daily mean temperature (°C), averaged with a moving window of ten days, under RCP 4.5 (in blue) and RCP 8.5 (in red) scenarios from 2030 to 2100. (Panel A) 2090-2099, (Panel B) 2080-2089, (Panel C) 2070-2079, (Panel D) 2060-2069, (Panel E) 2050-2059, (Panel F) 2040-2049, (Panel E) 2030-2039. Empty blue and red rectangles in each year represent the time from pit hardening *t*_0_ to harvest *t_H_* for RCP 4.5 and RCP 8.5, respectively and the ‘x’ symbols represent blooming failures.

**Figure S7:**
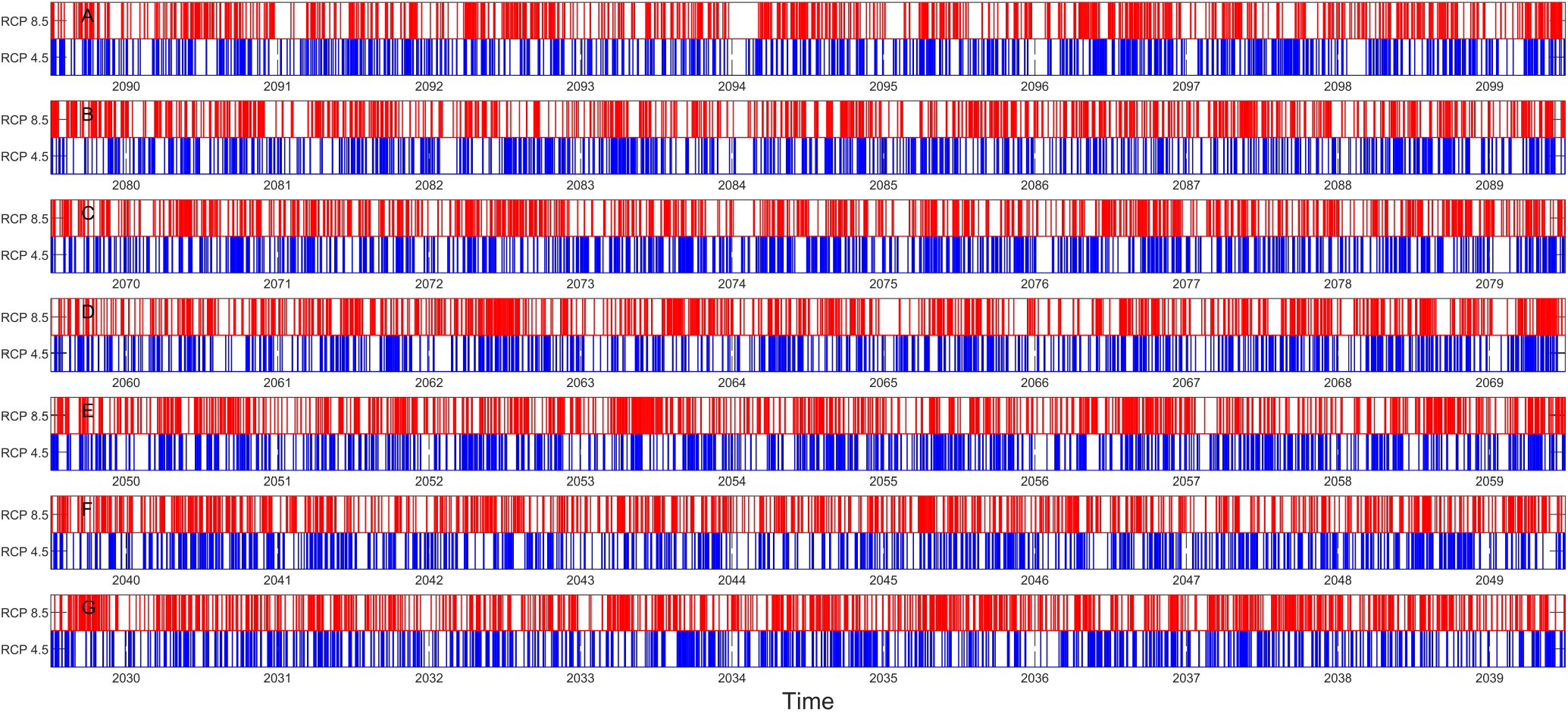
Daily precipitation occurrence under RCP 4.5 (in blue) and RCP 8.5 (in red) scenarios from 2030 to 2100. (Panel A) 2090-2099, (Panel B) 2080-2089, (Panel C) 2070-2079, (Panel D) 2060-2069, (Panel E) 2050-2059, (Panel F) 2040-2049, (Panel E) 2030-2039.

### Sub-models of fruit growth, abscission and fungal spore survival

#### Fruit growth model

Accordingly to [1], we described fruit growth by a logistic curve

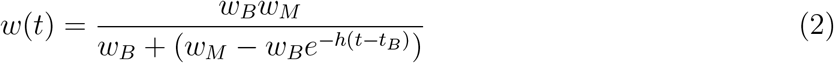

where *t_B_* is the time at bloom, which has been observed for both years (*t*_*B*,2014_=59 DOY and *t*_*B*,2015_=74 DOY); *w_B_* is the fruit weight at bloom; *w_M_* is the maximum fruit weight and *h* is the conversion rate of resources into fruit mass. We calibrated parameters *w_B_*, *w_M_* and h by minimizing the Sum of Square Errors *SSE*, and we retrieved the parameter distribution by the mean of the block bootstrap technique (Kreiss and Lahiri 2012),

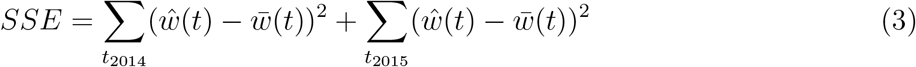

where 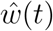 is the estimated fruit fresh weight and 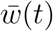 is the observed fruit fresh weight). The resulting estimated parameter values and 90 % CI are respectively *w_B_* 0.49 [0.39;0.61] g, *w_M_* 214 [198;233] g and *h* 5.6×10^−2^ [5.3;5.9]×10^−2^. Observed data and the simulated fruit growth curve are reported in Figure S8.

**Figure S8:**
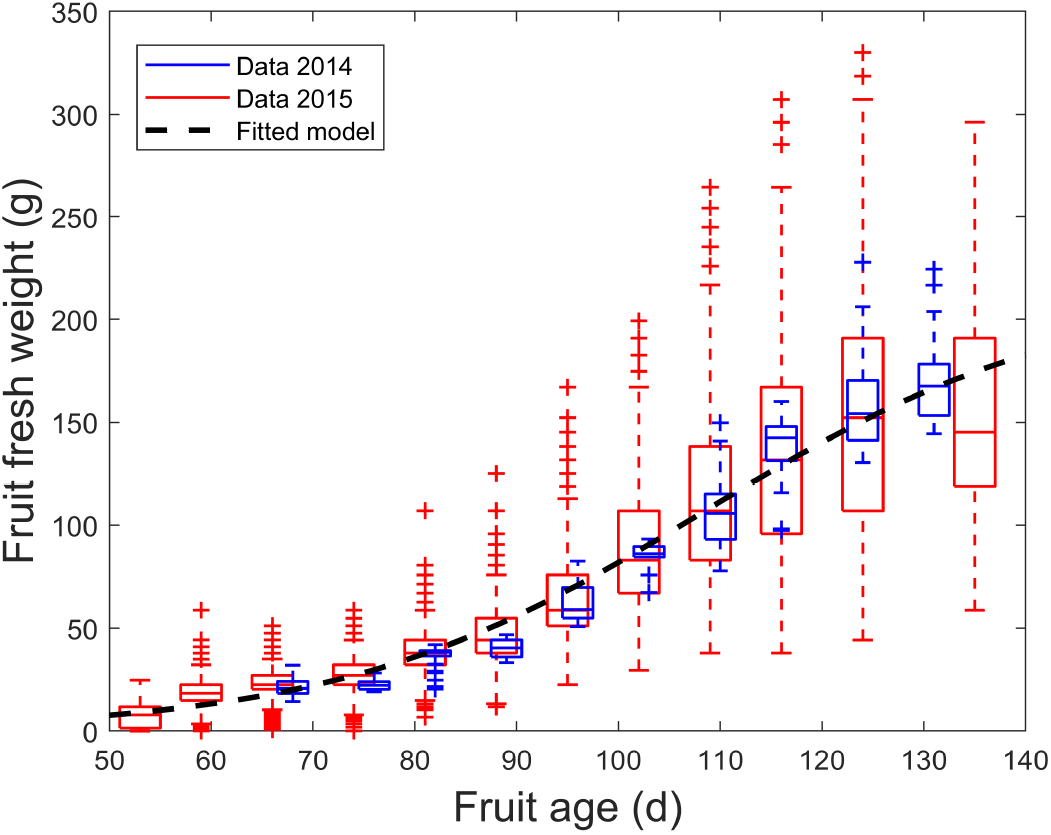
Fruit fresh weight vs. fruit age, calculated as days after bloom. Blue and red boxplots represent observed data of fruit fresh weight (g) in 2014 and 2015, respectively. Logistic fitted model is described by the black dashed line.

#### Fruit abscission model

We tested if the abscission rate *μ*(*t*), estimated from the data (see Table S1), varies either with the fruit density *N* (fruit m^−2^) and/or the fruit age *A* (days after bloom) (Figure S2). Fruit age may cause two major abscission events in stone fruits. The first post-blooming abscission that happens in the two weeks that follow the blooming is not studied in our work, because it does not occur in a period of brown rot spreading. The second abscission event, named “June drop”, takes place before the exponential weight growth of fruits, when the plant needs to conserve the necessary resources for this rapid growth [4]. This latter abscission event is the target of our model.

The two years 2014 and 2015 are characterized by different agronomic and climate conditions and, consequently, fruit densities and blooming times are different. We evaluated multiple competing hypotheses to test for the possible dependency of the abscission rate on fruit density *N*(*t*) and age *A*. We hypothesized that the abscission rate *μ*(*t*) could have a power dependency on fruit density *N*, and parabolic dependency on fruit age A. To test these alternative hypotheses, we incorporated them into 7 different models, which we compared to one another by evaluating the Akaike Information Criterion (Table S2 and main text). Competing hypotheses of fruit abscission rate dependency on fruit age and fruit density are incorporated them into 7 different models, which we compared to one another by evaluating the Akaike Information Criterion in Table S2.

**Table S2:** Ranking of the competing models listed here in increasing order of Akaike Information Criterion *AIC_c_*, reported together with their Root Mean Square Error (RMSE), i.e. 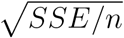

| Tested model | Model equation | $k$ | $\sqrt{SSE/n}$ | $AIC_c$ | $\Delta AIC_c$ |
| --- | --- | --- | --- | --- | --- |
| M6 | $\mu = (gA^2 + dA)N^b$ | 3 | $4.4 \times 10^{-3}$ | -165.7 | 0 |
| M5 | $\mu = \mu_0 N^b$ | 2 | $5.2 \times 10^{-3}$ | -163.7 | 2.0 |
| M7 | $\mu = (\mu_0 + gA^2 + dA)N^b$ | 4 | $4.4 \times 10^{-3}$ | -162.0 | 3.7 |
| M2 | $\mu = gA^2 + dA$ | 2 | $5.7 \times 10^{-3}$ | -160.5 | 5.1 |
| M4 | $\mu = \mu_0 + gA^2 + dA$ | 3 | $5.7 \times 10^{-3}$ | -157.5 | 8.2 |
| M1 | $\mu = \mu_0$ | 1 | $7.4 \times 10^{-3}$ | -154.7 | 11.0 |
| M3 | $\mu = N^b$ | 1 | $1.4 \times 10^{-2}$ | -133.6 | 32.1 |

Model M6 is selected as best model (Table S2). As shown by the model equations in Table S2, it describes the dependency of the abscission rate *μ* on fruit density according to a power law function and the dependency of *μ* on fruit age by a parabolic relationship characterized by a null intercept (Figure S9). The calibrated parameters values, together with their 90% CI, are the following: *b*=0.49 [−1.9× 10^−2^; 0.74], *d*= 9.1×10^−5^ [2.5;86] × 10^−5^ d^−1^, *g*=-6.8×10^−7^ [−81;-0.38]× 10^−7^ d^−2^.

**Figure S9:**
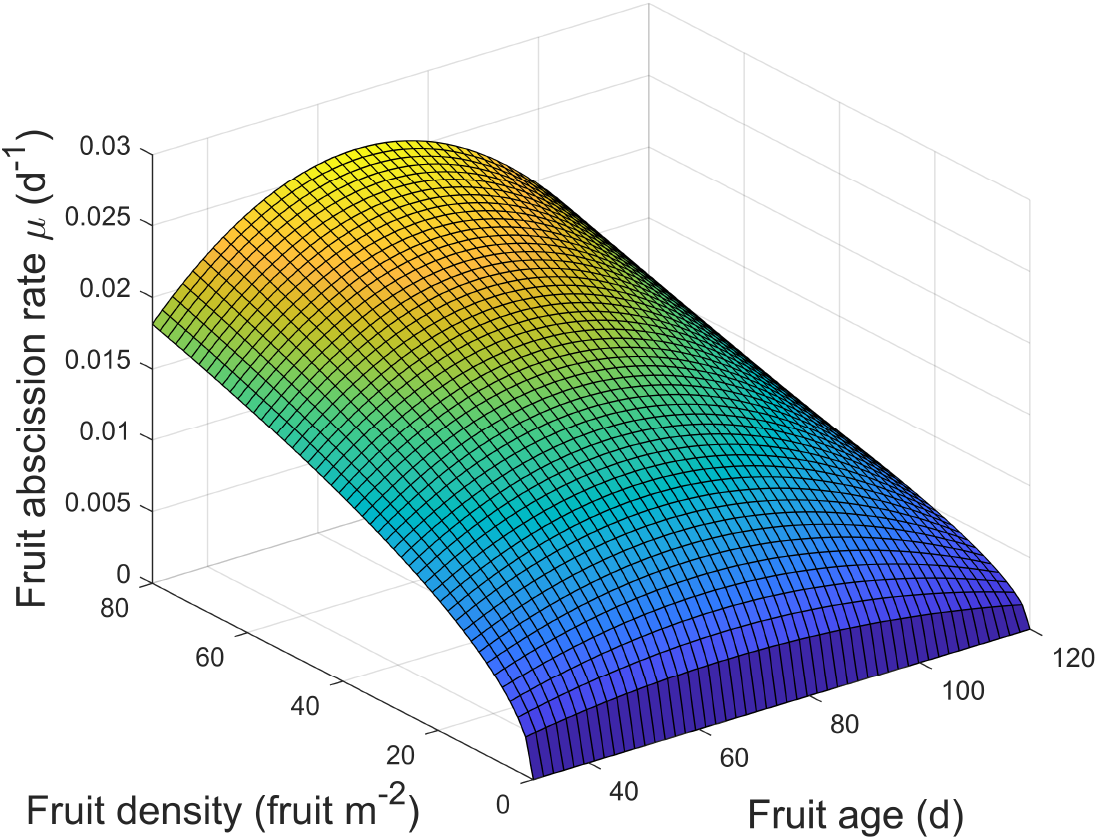
Functional representation of the fruit abscission rate dependency on fruit density (fruit m^−2^) and fruit age (days after bloom), as best estimated by the selected model M6 (Table S2)

#### Spore viability model

The parameter *η* represents the rate at which the spores deposited on fruits die or are wiped out so that the fruit can return to the susceptible state. It corresponds to the inverse of the mean duration of spore viability on the fruit surface. As spore lifespan can vary with the air temperature [3], we used conditions outlined in [7] to assess possible dependency of *η* from temperature *T*. According to Brown et al.’s (2004) metabolic theory of ecology, the influence of temperature in most ecological processes, such as death, can be modelled by the Van’t Hoff-Arrhenius equation of the form

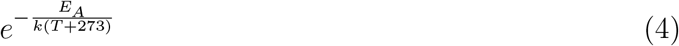

where *E_A_* is the activation energy, *k* is Boltzmann’s constant and *T* is temperature in °C. Therefore, we hypothesized a spore death, that is constant (M1) or temperature dependent (M2) according to Van’t Hoff-Arrhenius equation. We calibrated both models against data and selected the best one according to *AIC_c_* (see Table). Eventually, we assessed empirical parameter distribution via moving block bootstrap technique (1000 iterations).

The two competing evaluated models are reported in Table S3, together with model selection metrics.

**Table S3:** Akaike model selection and ranking

| Tested model | $k$ | $\sqrt{SSE/n}$ | $AIC_c$ | $\Delta AIC_c$ |
| --- | --- | --- | --- | --- |
| M2 | 2 | $8.3 \times 10^{-2}$ | -54.4 | 0 |
| M1 | 1 | 0.12 | -48.4 | 6.0 |

The selected best model (*M*2) accounts for temperature dependency according to metabolic theory. Its parametrization results to be *η*_0_=5.7×10^19^ [1.1× 10^16^;1.4× 10^26^] (d^−1^) and *r*=1.4×10^4^ [1.1;2.0] × 10^4^ (°C).

### Full epidemiological model

#### Conversion from minimum/maximum to average daily temperature

The minimum and the maximum temperature for fungal infection in plants have been estimated by [7] and [5]. They experimentally observed, in incubation chambers at controlled temperature, the range of temperatures at which infection is possible, as a consequence of spore development and germination. They found that temperature must remain between 3 °C and 30 °C (Tamm and Fluckiger 1993; Xu et al. 2001). In order to use only average daily temperatures as input of our epidemiological model, we assessed a relationship between the average daily temperatures, *T*, and the corresponding minimum and maximum daily temperatures, *T_min_* and *T_max_*, using our 2014 and 2015 data. Linear models (*T_min_* = *m_min_T* + *q_min_* and *T_max_* = *m_max_T* + *q_max_*, with *q_min_*=-2.6, *q_max_*=3.3, *m_min_*=0.86, *m_max_*=1.12) permitted to successfully relate average to minimum and maximum daily temperature (see Figure S10). We derived that the extreme minimum and maximum daily temperatures of 3 °C and 30 °C, in the considered region, correspond to the average daily temperature of 6.5°C and 23.3 °C, respectively. In regions with a different diurnal variation, one should reassess the relationship between average and extreme daily temperatures or opt for a direct use of minimal and maximal daily temperatures.

**Figure S10:**
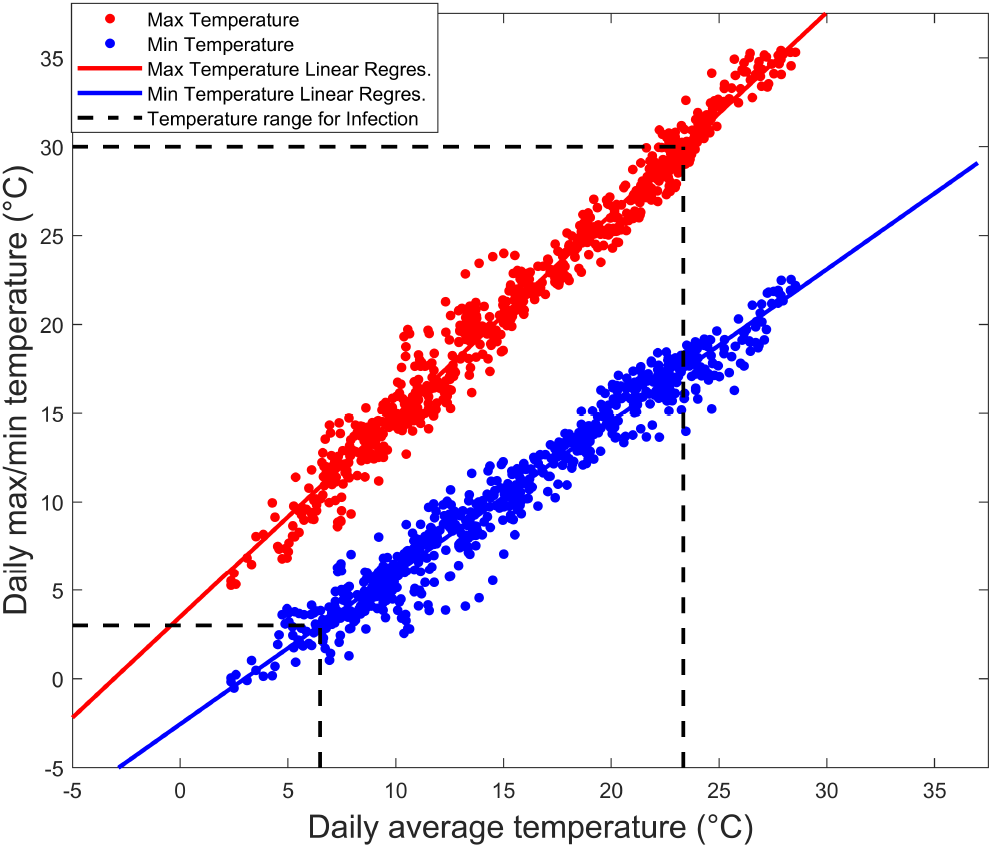
Regression between daily average temperature (°C) and minimum (in blue) and maximum (in red) temperatures (°C). Points represent data, lines are the regressions. Infection can happen in the temperature interval within the dashed black lines.

#### Akaike model selection

Competing epidemiological models and the alternative hypotheses underlying their formulations on possible parameter dependence on climatic conditions are reported in Table S4.

**Table S4:** Tested competing models for the exposure rate of S (λ) and the progression coefficient (*γ*) in Akaike model selection

| Model | $\lambda$ | $\gamma$ |
| --- | --- | --- |
| M1 | $\lambda_0$ const | $\gamma_0$ const |
| M2 | $\lambda_0$ const | $\gamma_0(T)$ |
| M3 | $\lambda_0$ const | $\gamma_0(P)$ |
| M4 | $\lambda_0$ const | $\gamma_0(T, P)$ |
| M5 | $\lambda(P)$ | $\gamma_0$ const |
| M6 | $\lambda(P)$ | $\gamma_0(T)$ |
| M7 | $\lambda(P)$ | $\gamma_0(P)$ |
| M8 | $\lambda(P)$ | $\gamma_0(T, P)$ |

The results of the Akaike model selection is reported in Table S5.

**Table S5:**
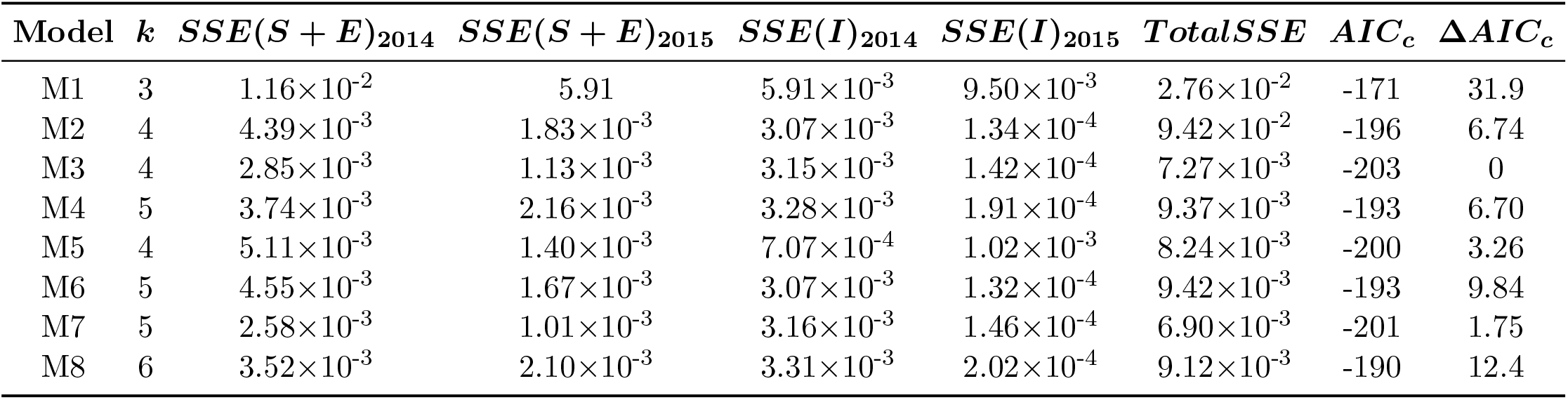
Akaike model selection. Sum Square Error *SSE* of Susceptible and Exposed fruit (*S*+*E*) and of Infectious fruits (*I*) for 2014, 2015 and total are reported. Corrected Akaike Information Criterion *AIC_C_* and Δ*AIC_C_* are reported.

| Model | $k$ | $SSE(S + E)_{2014}$ | $SSE(S + E)_{2015}$ | $SSE(I)_{2014}$ | $SSE(I)_{2015}$ | $TotalSSE$ | $AIC_c$ | $\Delta AIC_c$ |
| --- | --- | --- | --- | --- | --- | --- | --- | --- |
| M1 | 3 | $1.16 \times 10^{-2}$ | 5.91 | $5.91 \times 10^{-3}$ | $9.50 \times 10^{-3}$ | $2.76 \times 10^{-2}$ | -171 | 31.9 |
| M2 | 4 | $4.39 \times 10^{-3}$ | $1.83 \times 10^{-3}$ | $3.07 \times 10^{-3}$ | $1.34 \times 10^{-4}$ | $9.42 \times 10^{-2}$ | -196 | 6.74 |
| M3 | 4 | $2.85 \times 10^{-3}$ | $1.13 \times 10^{-3}$ | $3.15 \times 10^{-3}$ | $1.42 \times 10^{-4}$ | $7.27 \times 10^{-3}$ | -203 | 0 |
| M4 | 5 | $3.74 \times 10^{-3}$ | $2.16 \times 10^{-3}$ | $3.28 \times 10^{-3}$ | $1.91 \times 10^{-4}$ | $9.37 \times 10^{-3}$ | -193 | 6.70 |
| M5 | 4 | $5.11 \times 10^{-3}$ | $1.40 \times 10^{-3}$ | $7.07 \times 10^{-4}$ | $1.02 \times 10^{-3}$ | $8.24 \times 10^{-3}$ | -200 | 3.26 |
| M6 | 5 | $4.55 \times 10^{-3}$ | $1.67 \times 10^{-3}$ | $3.07 \times 10^{-3}$ | $1.32 \times 10^{-4}$ | $9.42 \times 10^{-3}$ | -193 | 9.84 |
| M7 | 5 | $2.58 \times 10^{-3}$ | $1.01 \times 10^{-3}$ | $3.16 \times 10^{-3}$ | $1.46 \times 10^{-4}$ | $6.90 \times 10^{-3}$ | -201 | 1.75 |
| M8 | 6 | $3.52 \times 10^{-3}$ | $2.10 \times 10^{-3}$ | $3.31 \times 10^{-3}$ | $2.02 \times 10^{-4}$ | $9.12 \times 10^{-3}$ | -190 | 12.4 |

The fitting of the model that is climate independent (M0), where spore death rate *η*, infection rate *σ* and exposure rate λ are temperature and precipitation independent, is reported in Figure S11B. Note in particular the mismatch between simulation and data for the infectious compartment in 2015.

**Figure S11:**
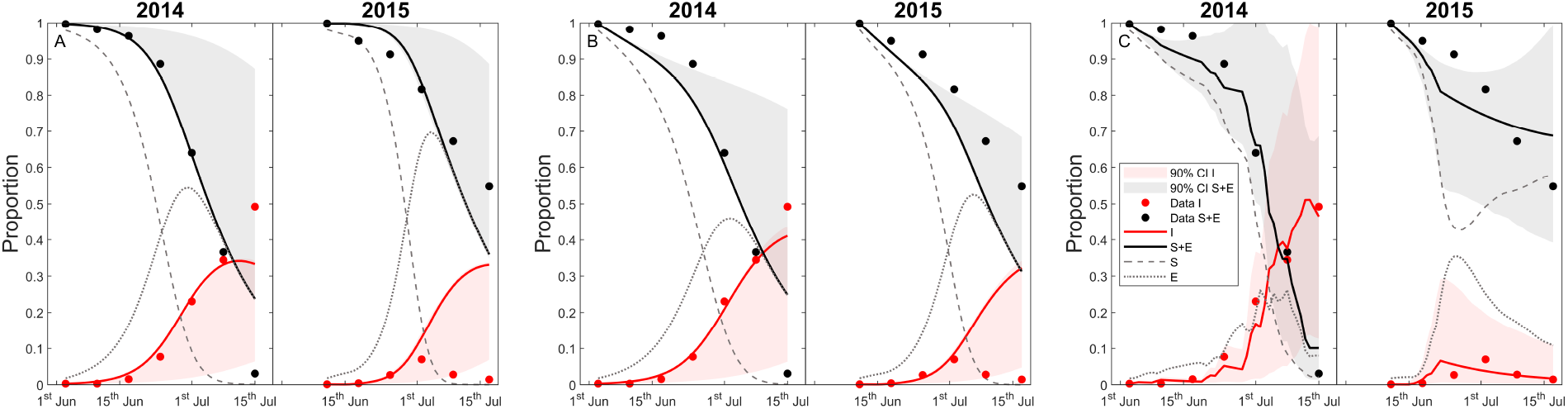
Modeling outline step by step: temporal dynamics of Susceptible (*S*), Exposed (*E*), Infected (*I*), and Symptomless (*S* + *E*) fruits, in 2014 and 2015. In panel A, the original model Bevacqua et al. [1] calibrated on the two years (*SSE*=2.82×10^−2^, see eq 10 main text). In panel B, the original model enhanced with with fruit abscission (*SSE*=2.76×10^−2^, M0 in Table S5). In panel C, the original model enhanced with fruit abscission, temperature dependence of parameter *η* and precipitation dependence of (*σ*) (SSE=7.27×10^−3^, M3 in Table S5). Lines represent simulations and symbols represent observed values (black circles for symptomless and red circle for infectious fruit abundance. Shaded areas (same color coding) indicate the predicted 90% confidence bands, obtained via bootstrap technique, for symptomless and infectious fruits, respectively.

**Figure S12:**
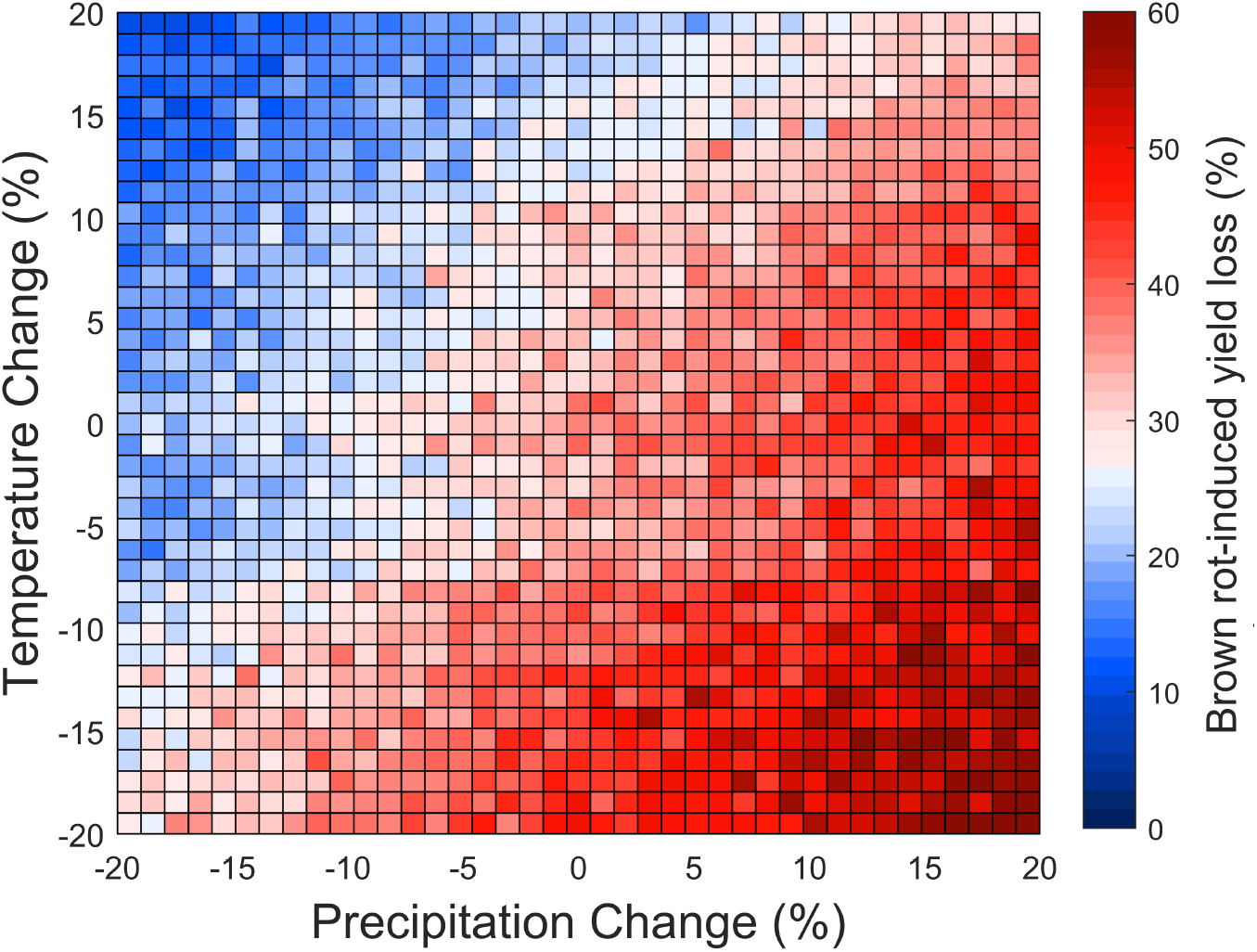
Predicted percentage of yield losses due to brown rot of a virtual untreated peach orchard under different scenarios of temperature and rain occurrence, that have been varied by +/-20 % compared to the average conditions of 2010-2019.

